# Exploring the vomeronasal organ in an endangered antelope species

**DOI:** 10.1101/2023.03.09.531847

**Authors:** Mateo V. Torres, Irene Ortiz-Leal, Andrea Ferreiro, José Luis Rois, Pablo Sanchez-Quinteiro

## Abstract

The dama gazelle is a threatened and scarcely studied species of northern Africa. Human pressure has depleted the population of dama gazelles from tens of thousands to a few hundred individuals. Since 1970, after deriving from a founder population of the last 17 surviving individuals in the Western Sahara, it has been reproduced naturally in captivity. Although certain aspects of the reproductive biology of the dama gazelle have been established in preparation for the future implementation of assisted reproductive technology there is a lack of information regarding the role of semiochemical-mediated communication in the sexual behavior of the dama gazelle. This is partially due to the lack of a neuroanatomical and morphofunctional characterization of the dama gazelle vomeronasal organ (VNO); the sensory organ responsible for the processing of pheromones. This study aims to determine the presence in the dama gazelle of a VNO fully equipped to carry out its neurosensory function and to contribute to the understanding of the interspecific variability of the VNO of ruminants. Employing histological, lectin-histochemical and immunohistochemical techniques we have performed a detailed morphofunctional evaluation of the dama gazelle VNO along its entire longitudinal axis. The findings suggest that studies of the VNO should take a similar approach, as there are significant structural and neurochemical transformations that the organ exhibits as a whole. This study contributes to the understanding of the VNO in dama gazelles and provides a basis for future studies on semiochemical-mediated communication and reproductive management of this species.

## 1. INTRODUCTION

The dama gazelle (*Nanger dama*) is one of the most singular, threatened and scarcely studied gazelle species of northern Africa (Abáigar et al., 2019). It is the tallest and heaviest of all gazelles, with males weighing up to 70 kg. There are unique physiological, ecological, and behavioral adaptations that allow this species to thrive in the arid and semi-arid environments of North Africa, where it is a member of the group of organisms known as Sahelo-Saharian Ungulates (SSU) (Durant et al., 2014). According to their geographic distribution and coat color, three subspecies can be distinguished: addra gazelle (*Nanger dama ruficollis*), mhorr gazelle (*Nanger dama mhorr*) and dama gazelle (*Nanger dama dama*) (Scholte, 2019). Their ruminant status is reflected in their diet, which is based on foraging on grasses and browsing on shrubs or acacias. They have a nomadic behavior and live in herds composed of a single dominant male and several females.

Human pressure has depleted the population of dama gazelles and caused a disruption in their ecological system. Since the 1950s, there has been a drastic decline in the dama gazelle population from tens of thousands to a few hundred individuals. This phenomenon was attributed to a combination of two factors. First, habitat degradation, largely due to increased grazing by domestic livestock (Devillers et al., 2005). And secondly, to intense hunting activity (Grettenberger & Newby, 1986; Cloudsley-Thompson, 1992). Currently, the dama gazelle is considered a critically endangered species according to the red list of the International Union for Conservation of Nature (IUCN). Since 1970, after deriving from a founder population of the last 17 surviving individuals in the Western Sahara (Alados et al., 1988), it has been reproduced naturally in captivity (Holt et al., 1996). The remaining populations are all very small and extremely fragmented (Lamarque et al., 2007). Looking to the future, the situation of the dama gazelle is not expected to improve. On the one hand, it is considered that the quality of its habitat will continue to deteriorate (Bronson, 2009; Durant et al., 2014). On the other hand, the hunting conflict that severely undermines its population remains unresolved. Additionally, frequent fights between males have been reported in this species when reintroducing populations into their natural habitat. This is a problem that is difficult to solve, since the clashes are due to the limited space available (Cassinello & Pieters, 2000).

Maintaining a sustainable population of endangered wildlife species is a challenge, which in the case of the dama gazelle is made even more difficult by the aforementioned aspects. Because reproduction is fundamental to species survival, understanding reproductive mechanisms is a high priority. It is well established that there are enormous differences in the specifics of how each species reproduces, even those in the same phylogenetic clade (i.e., family; Wildt et al., 2001). A recent survey of 50 species concluded that there are as many mechanistic variations in reproduction as there are species (Wildt et al., 2010). Although certain aspects of the reproductive biology of the dama gazelle have been established in preparation for the future implementation of assisted reproductive technology (Pickard et al., 2001, 2003; Wojtusik et al., 2017; Arregui et al., 2021), there is, to our knowledge, no information regarding the role of semiochemical-mediated communication in the sexual behavior and reproductive physiology of the dama gazelle. Sexual attraction, mother-young interactions, estrus indication, induction and synchronization, puberty acceleration, reducing the postpartum anestrus, hormonal stimulation, and enhancing the penis erection are some of the classical events influenced by pheromones in ruminants (Archunan et al., 2014). Moreover, pheromones are also strongly involved in the control of stress regulation mechanisms in mammals. Therefore, they could be used at the time of reintroduction of specimens in order to appease the aggressiveness of males (Archer et al., 2022).

Although there is plenty of scope for the application of cattle pheromones in the reproduction and management of the dama gazelle, this requires the previous neuroanatomical and morphofunctional characterization of its vomeronasal organ (VNO), the sensory organ responsible for the processing of semiochemicals as pheromones (Powers & Winans, 1975; Kunkhyen et al., 2017), kairomones (Isogai et al., 2011; Fortes-Marco et al., 2013), and molecules of the major histocompatibility complex (Leinders-Zufall et al., 2000, 2014). The organ consists of two tubular, blind-ending, and elongated structures located at the base of the nasal septum (Poran, 1998). Inside them is the vomeronasal duct, which establishes a direct communication with the nasopalatine duct and through it with the external environment (Takami, 2002). The vomeronasal duct is lined by vomeronasal epithelium, which is composed of pheromone-sensing receptor neurons (Eisthen, 1992; Zufall et al., 2002). These molecules, coming from the nasal or oral cavities, enter the VNO dissolved in mucus and activate the vomeronasal neuroepithelium (Sankarganesh et al., 2022). Transduction takes place in its neuroreceptors and is conveyed by the vomeronasal nerves to the accessory olfactory bulb, the first integrating center for pheromonal information in the central nervous system (Halpern, 1987; Mori, 1993; Villamayor et al., 2020; Ortiz-Leal, Torres, Villamayor, et al., 2022). The morphological and immunohistochemical study of the VNO of the dama gazelle will answer questions such as the existence of a functional communication between its VNO and the environment, or whether its VNO is fully equipped to carry out its neurosensory function, having a complete external envelope, a differentiated neuroepithelium, and a parenchyma with the necessary glandular, vascular and nervous elements (Salazar, Sánchez Quinteiro, et al., 1997; Salazar & Quinteiro, 1998). Moreover, this study will contribute to improve the understanding of the basis of the interspecific variability of the VNO of ruminants and mammals in general.

The dama gazelle belongs to the order of the Artiodactyla, suborder Ruminantia, and infraorder Pecora. Within the latter, it is part of the Bovidae family, the one with the largest number of species. Other families in this group are the Antilocapridae (pronghorns), Giraffidae, Moschidae (musk deers), and Cervidae. Available information on the VNO in these families is limited to anatomical and histological studies of the giraffe VNO (Kondoh, Nakamura, et al., 2017; Hart & Hart, 2023), the specific study of glyconjugates in the musk deer organ (Kondoh et al., 2020), whereas the Cervidae family has been studied more extensively. Both the reindeer (*Rangifer tarandus*) (Bertmar, 1981), elk (*Alces alces*) (Vedin et al., 2010), sika deer (*Cervus nippon*) (Matsubara et al., 2019) and Korean roe deer (*Capreolus pygargus*) (Park et al., 2014; Shin et al., 2017) showed a functional and well-developed VNO.

Although the study of VNO in bovids has been more extensive, it has mainly focused on domestic species such as cow (Jacobs et al., 1981; Taniguchi & Mikami, 1985; Adams, 1986; Salazar et al., 2008; Jang et al., 2021), goat (Ladewig & Hart, 1980; Park et al., 2013; Yang et al., 2021) and sheep (Kratzing, 1971; Salazar et al., 1998, 2000; Salazar, Lombardero, Alemañ, et al., 2003; Salazar et al., 2007; Ibrahim et al., 2013; Barrios et al., 2014; Ibrahim, 2018). Little research has been done, however, on the VNO of wild bovids. Noteworthy are the substantial differences discovered in the investigation of alcelaphine antelopes, which showed that three of the species studied, *Damaliscus iunatus*, *Alcelaphus buscelaphus*, and *Connochaetes taurinus*, either lacked the flehmen response or had no direct communication between the oral cavity and the VNO via the incisive duct. The general histology study showed however an apparently developed VNO (Hart et al., 1988). These striking anatomical observations found their genetic correspondence in a recent study of ancV1R gene expression (Zhang & Nikaido, 2020), considered a reliable diagnostic indicator of VNO function. The authors observed the inactivation of ancV1Rs expression in seals, sea otter, giant otter, fossa, the owl monkey, and alcelaphine antelopes. Beyond antelopes, information on wild ruminants is limited to the morphofunctional study of the VNO of the water buffalo (*Bubalus bubalis*) (Emam et al., 2016), the study of the glandular secretion in *Gazella subgutturosa* (Abood & Hussein, 2018) and the CT study of the nasal cavity of the saiga antelope (*Saiga tatarica*) which revealed the presence of the VNO in this species (Clifford & Witmer, 2004). Despite the limited information available, these observations point to the presence of a wide morphological diversity in the family Bovidae. Once again, the interspecies extrapolation of information relative to the anatomy and histology of the VNS can be difficult and risky (Salazar et al., 2016).

In this study of the VNO of the dama gazelle, we set a double objective: to define the morphological and immunohistochemical basis of the VNO of the dama gazelle, as a first necessary step to carry out rigorously the implementation of chemical communication in the reproductive management of the species, and secondly to contribute to fill the gaps in the knowledge of the VNO of the family Bovidae.

To this end, we have carried out a comprehensive morphofunctional study of the dama gazelle OVN using histological, histochemical and immunohistochemical techniques. As a remarkable feature in our approach, we have performed an exhaustive study along the entire longitudinal axis of the OVN evaluating the existence of differences in terms of its microscopic structure, but also in terms of the immunohistochemical labeling pattern. The wide range of differences found suggests that studies of the VNO should take a similar approach, highlighting the important structural and neurochemical transformations that the organ exhibits as a whole.

## 2. MATERIAL & METHODS

### 2.1. Samples

To carry out our study, we had a healthy male adult dama gazelle (*Nanger dama*), which was donated by the Natural Park “Marcelle Natureza” (Outeiro de Rei, Spain) due to its death by accidental causes, specifically a neck fracture when the animal was engaged in social play. Less than 12 hours after the animal’s death, the head was separated from the carcass and the skin, tongue, jaws, teeth, nose and eyes were removed. A cross-section was made with the aid of a rotary saw to separate the snout, where the VNOs are located, from the rest of the skull. In the snout sample, the bony sides of the nasal cavity and the nasal turbinates were removed. This allowed to identify the location of the VNOs on both sides of the basal part of the nasal septum. Subsequently, a transverse cut was made to divide the nasal septum centrally to facilitate the entry of the fixative into the vomeronasal ducts. The resulting samples were immersed in freshly prepared Bouin’s fluid and, after 24 hours, transferred to 70% ethanol. The following day, the VNO were dissected, and multiple cross-sections were made equidistant along their entire length in order to analyse the characteristics of each area. This procedure was carried out under a Zeiss OPMI 1 surgical microscope. The bone tissue surrounding the VNO ventrally and medially was dissected from all samples, with the exception of those at the most rostral levels. These rostral pieces were decalcified for a week to microscopically study the topographic relationship of the VNO with the incisive duct. The decalcifying agent used was Shandon TBD-1 Decalcifier (Thermo, Pittsburgh, PA), and it was applied while stirring continuously at room temperature. All samples were marked and embedded in paraffin, thus obtaining the corresponding blocks. They were cut with a Leica Reichert Jung microtome with a thickness of 5–7 μm, depending on the tissue to be processed.

### 2.2 Histological staining

In order to be able to visualize the different components of the analyzed tissue, we used hematoxylin-eosin (HE) as general staining. Additionally, periodic acid Schiff (PAS) and alcian blue (AB) stains were used to identify neutral and acidic mucopolysaccharides respectively.

### 2.3 Lectin histochemical staining

Lectins, naturally occurring carbohydrate-binding molecules, can be used to detect glycoconjugates associated with certain cell populations in the tissues tested. For this reason, we have used the following five lectins in our research:

- UEA (Vector-L1060), which comes from gorse, *Ulex europaeus*, and marks the L-fucose pathway (Kondoh et al., 2018).
- LEA (Vector-B1175), which comes from tomato, *Lycopersicon esculentum*, and recognizes N-acetyl-glucosamine (Salazar & Sanchez-Quinteiro, 2003).
- VVA, which is isolated from fodder vetch, *Vicia villosa*, binds specifically to N-acetylgalactosamine structures (Tomiyasu et al., 2018).
- SBA (Vector-B1235), which comes from fodder soybean, *Glycine max*, binds preferentially to oligosaccharide structures with N-acetylgalactosamine and to galactose residues (Park et al., 2012).
- STA (Vector-B1165), which is isolated from potato, *Solanum tuberosum*, binds oligomers of N-acetylglucosamine and some oligosaccharides containing N-acetylglucosamine and N-acetylmuramic acid (Chun et al., 2023).

The UEA lectin was employed unmodified and identified with an anti-UEA antibody, whereas the other four lectins, VVA, STL, LEA, and SBA, were biotinylated. The following procedures were followed:

#### 2.3.1 Biotynilated lectins protocol

The lectin protocols employed are fully explained in Ortiz-Leal et al. (2022) (i) The tissular endogenous peroxidase activity was blocked to avoid interference with the final developing step by incubating in a 3% H2O2 solution for 10 minutes. (ii) Sections were incubated for 30 minutes at room temperature in 2% bovine serum albumin (BSA), which prevents nonspecific binding. (iii) The slides were incubated in a humidified chamber overnight at 4°C in the biotinylated lectin diluted in 0.5% BSA at 1:200. The following day, samples were incubated (iv) for 90 min at room temperature in Vectastain ABC complex (Vector Laboratories, Burlingame, CA, USA). Finally, (v) the sections were developed by incubating the sections in a solution of 0.05% diaminobenzidine (DAB) and 0.003% H2O2 for 5-10 minutes. The reaction was monitored with the microscope.

#### 2.3.1 UEA lectin protocol

The protocol for UEA began with the same first two steps. Subsequently, (iii) incubation with UEA lectin (1:60 in 0.5% BSA) was carried out for 1 h at room temperature; (iv) 3 washes of 5 min in 0.1M phosphate buffer (PB, pH 7.2). (v) Afterwards the sections were incubated in a humidified chamber (overnight 4°C) in a peroxidase conjugated immunoglobulin against UEA (DAKO, P289; 1:50). Finally, the samples were developed (vi) by incubation in the same DAB solution as that employed in the byotinilated-lectins protocol. Controls were performed for both protocols, both without the addition of lectins and with the preabsorption of lectins, by using an excess amount of the corresponding sugar.

### 2.4 Immunohistochemical staining

Using immunohistochemistry, the gazella dama VNO was examined in-depth. The anti-Gαi2 and anti-Gαo antibodies are particularly useful because they label the transduction cascade for V1R and V2R vomeronasal receptors, respectively (Halpern et al., 1995). Antibodies were also used to detect olfactory marker protein (OMP), which is expressed in mature neurons in both olfactory subsystems (Rodewald et al., 2016). The calcium-binding proteins calbindin (CB), calretinin (CR), parvalbumin (PV), and secretagogin (SG) participate in the regulation of cytosolic free calcium ion concentrations in neurons. The distribution of proteins has previously been recognized as a useful neuronal marker for identifying specific brain regions and discrete neuronal subpopulations (Kishimoto et al., 1993; Coppola & Disney, 2018; Pérez-Revuelta et al., 2020; de Góis Morais et al., 2021). Cytokeratin 8 forms part of the intermediate filament cytoskeleton of epithelial cells. Accordingly, it is an effective marker for olfactory and vomeronasal sustentacular cells (Khan et al., 2021). Enolases are glycolytic enzymes that may be involved in cellular differentiation. γ enolase, specific of the nervous tissue, is a good marker for mature olfactory and vomeronasal neuroreceptor cells (Takahashi et al., 1984). PGP 9.5, also known as ubiquitin C-terminal hydrolase, is highly specific to neurons and to cells of the diffuse neuroendocrine system. In the VNO it identifies neuroreceptor cells but not supporting or basal cells (Taniguchi et al., 1993).

The protocol of the immunohistochemical labelling is explained in detail in Torres et al., 2021. Briefly, the initial step consisted of inhibiting endogenous peroxidase activity with 3% H2O2. Then (ii) non-specific binding was blocked for 30 minutes using 2.5% normal horse serum from the ImmPRESS Anti-Mouse IgG / Anti-rabbit IgG Reagent Kit (Vector Laboratories, CA, USA). (iii) Subsequently, the primary antibody was added to the corresponding dilution and incubated overnight (4°C). The following day, (iv) the samples were incubated at room temperature for 20 minutes with the corresponding ImmPRESS VR Polymer HRP anti-rabbit IgG reagent. After (v) rinsing in Tris buffer (0.2 M pH 7.61) for 10 minutes, (vi) the samples were developed using DAB as a chromogen in the same manner as for lectins.

All immunohistochemical protocols were run with the appropriate controls. In the absence of a positive control from dama gazelle, we reproduced the whole histochemical procedure with mouse tissues known to express the respective antigens. Samples for which the primary antibody was replaced by antibody diluent were used as negative controls.

### 2.5 Antibody characterization and specificity

Table 1 gives information on all antibodies, including their suppliers, dilutions, immunogen targets, and Research Resource Identifiers (RRID). In each instance, the immunostaining patterns produced by these antibodies in dama gazelles were comparable to those previously observed in a variety of mammalian species. References pertinent to each antibody are also listed in Table 1.

**Table 1.**
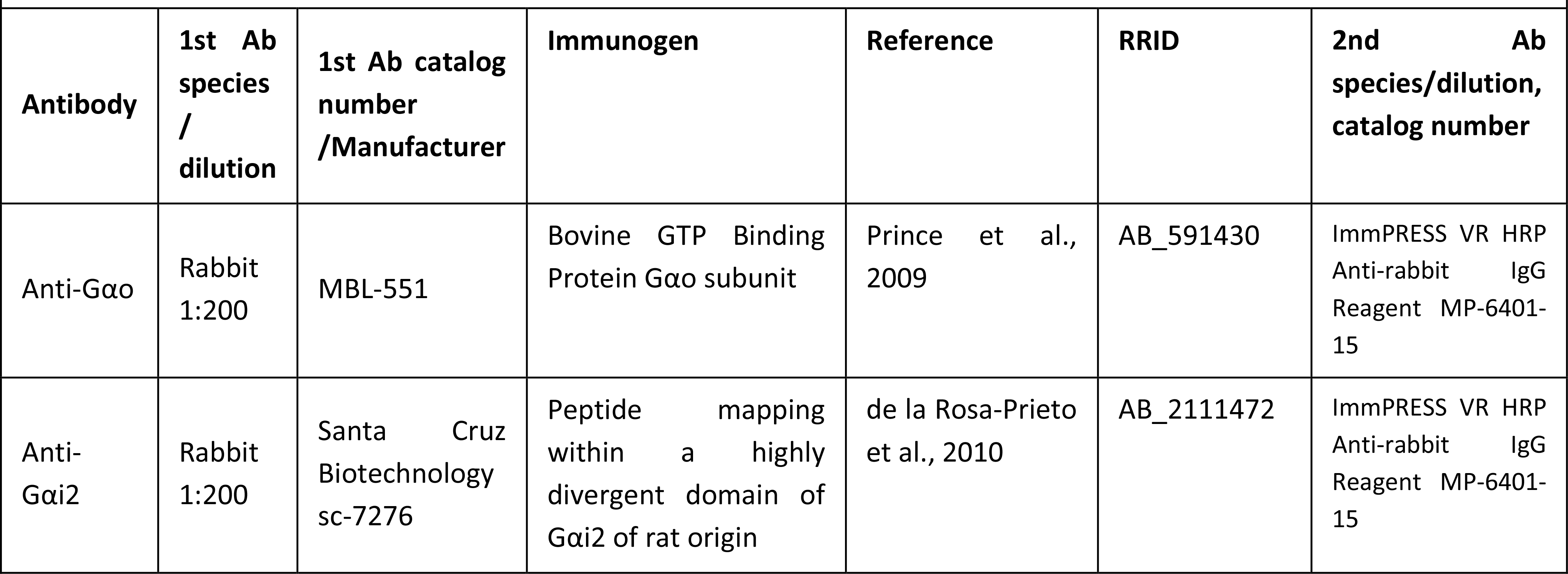
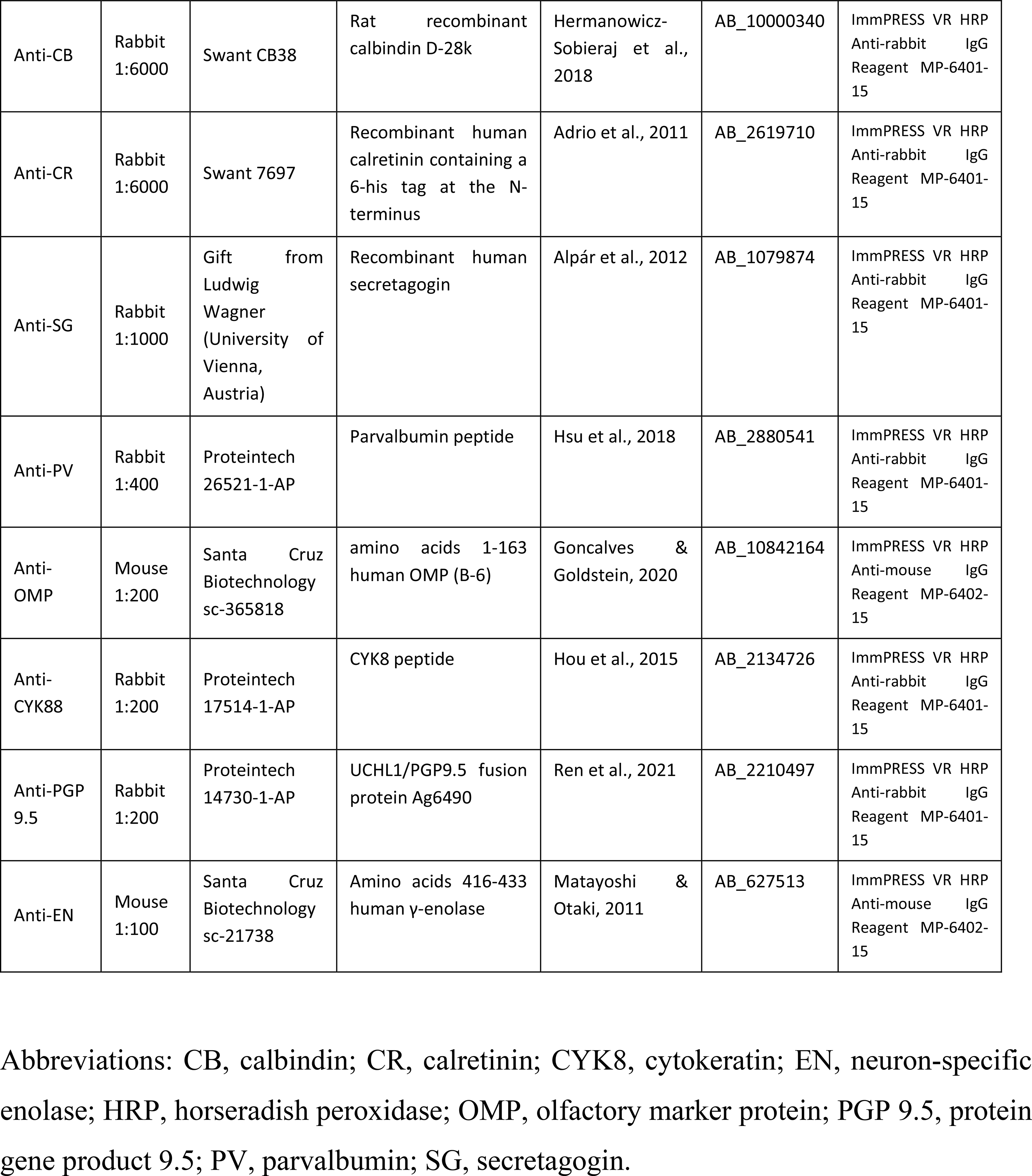
Detailed information on the antibodies used in this study: Species of elaboration, dilution, catalogue number, manufacturer, target immunogens, relevant reference for each antibody, RRID codes, and secondary antibody employed.

### 2.6 Acquisition of images

The images that make up the different illustrations were taken with a Olympus SC180 digital camera coupled to a Olympus BX50 microscope.

## 3. RESULTS

### 1. Macroscopic study

The vomeronasal organ of the dama gazelle consists of two tubular and elongated structures located at the base of the nasal septum, extending from the level of the incisive papilla to a level corresponding approximately to the rostral border of the lacrimal bone. To account for the structural and neurochemical changes that may occur along the longitudinal axis of such a lengthy structure, we conducted a histological serial study that resulted in the definition of fifteen transverse levels. These levels are depicted in Fig. 1A, which shows a lateral view of the head of the dama gazelle.

**Figure 1.**
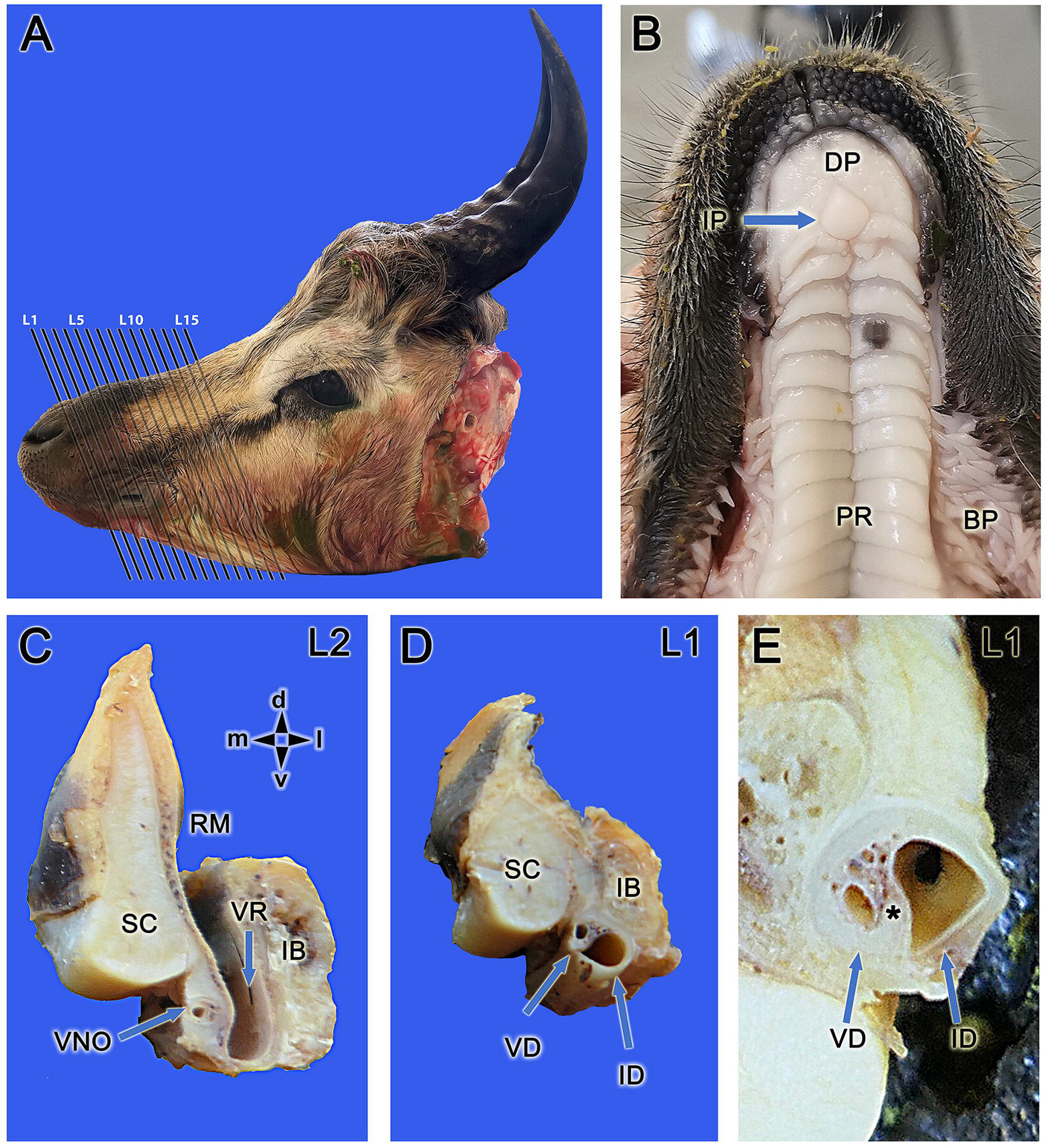
Dissection of the VNO of dama gazelle. A. Lateral view of the head showing the fifteen levels that correspond to the histological series shown in Figure 2. **B.** Ventral view of the palate, showing the opening of the incisive duct in the incisive papilla (IP). **C-D.** Sections of the decalcified nasal cavity, at the first two levels showed in **A**, to study the communication between the VNO and the external environment. The most rostral level (L1) is shown in **D**. The wide and lateral incisive duct (ID) runs parallel to the smaller vomeronasal duct (VD). The topographical relationship with the midline septal cartilage (SC) and the lateral incisive bone (IB) can also be seen. Both ducts are surrounded by a well-developed cartilaginous tissue, however, the communication between the VD and the ID has not yet been established. The next caudal level (L2) is shown in **C**. The cartilaginous envelope of the VNO is thick, and its dorsal process contacts the SC. At this stage, the incisive duct has opened in the ventral recess of the nasal cavity (VR). **E.** A higher magnification of the vomeronasal L1, shows how the cartilaginous envelope of both the VNO, and the ID is unique. The asterisk shows the development of the erectile tissue of the VNO, formed by numerous blood vessels, located between the VD and the ID. BP, buccal papillae; DP, dental pad; PR, palatal rugae; RM, respiratory mucosa. Orientation: d, dorsal; l, lateral; m, medial; v, ventral.

In artiodactyls, the communication between the vomeronasal duct and the external environment is made possible in an indirect manner by the opening of the vomeronasal duct into the incisive duct, which is also known as the nasopalatine duct. The incisive duct usually opens into the oral cavity via the incisive papilla, and it extends into the nasal cavity by its entrance into an aperture in the ventral recess of the nasal cavity. Since the presence of an incisive papilla is not a characteristic shared by all species of the family Bovidae studied to date, we paid particular attention in the dama gazelle to the study of the presence of an incisive papilla and the demonstration of a functional communication between the vomeronasal duct and the incisive duct. The examination of the hard palate of the dama gazelle (Fig. 1B) demonstrated the opening of the incisive duct in a well-defined incisive papilla.

It was feasible to examine the topographic relationships between the vomeronasal and incisive ducts, as well as the septal cartilage of the nasal cavity and the incisive bone, thanks to the decalcification of the two rostral blocks (L1-L2) (Fig. 1C,D). Both ducts run in parallel to one another while occupying the region that corresponds to the palatal fissure, with the ID duct being considerably larger in caliber than the other duct. It is noticeable how both passages, despite being separated by a broad band of erectile tissue, are encased in a common cartilaginous envelope (Fig. 1E). At level L2, the incisive duct has already opened to the ventral recess of the nasal cavity, while the vomeronasal duct exhibits a large cartilaginous envelope (Fig. 1C). The performed sections did not permit macroscopic identification of the communication between both ducts; hence, this issue could not be clarified until the entire histologic series of the VNO was performed.

### 2. Microscopic study

#### 2.1. Histological study

The histological series of the VNO comprises 15 levels (Fig. 2) and allows to discriminate the changes in the shape and dimensions of its main three components: the vomeronasal cartilage that encapsulates the vomeronasal duct and the parenchyma between them.

**Figure 2.**
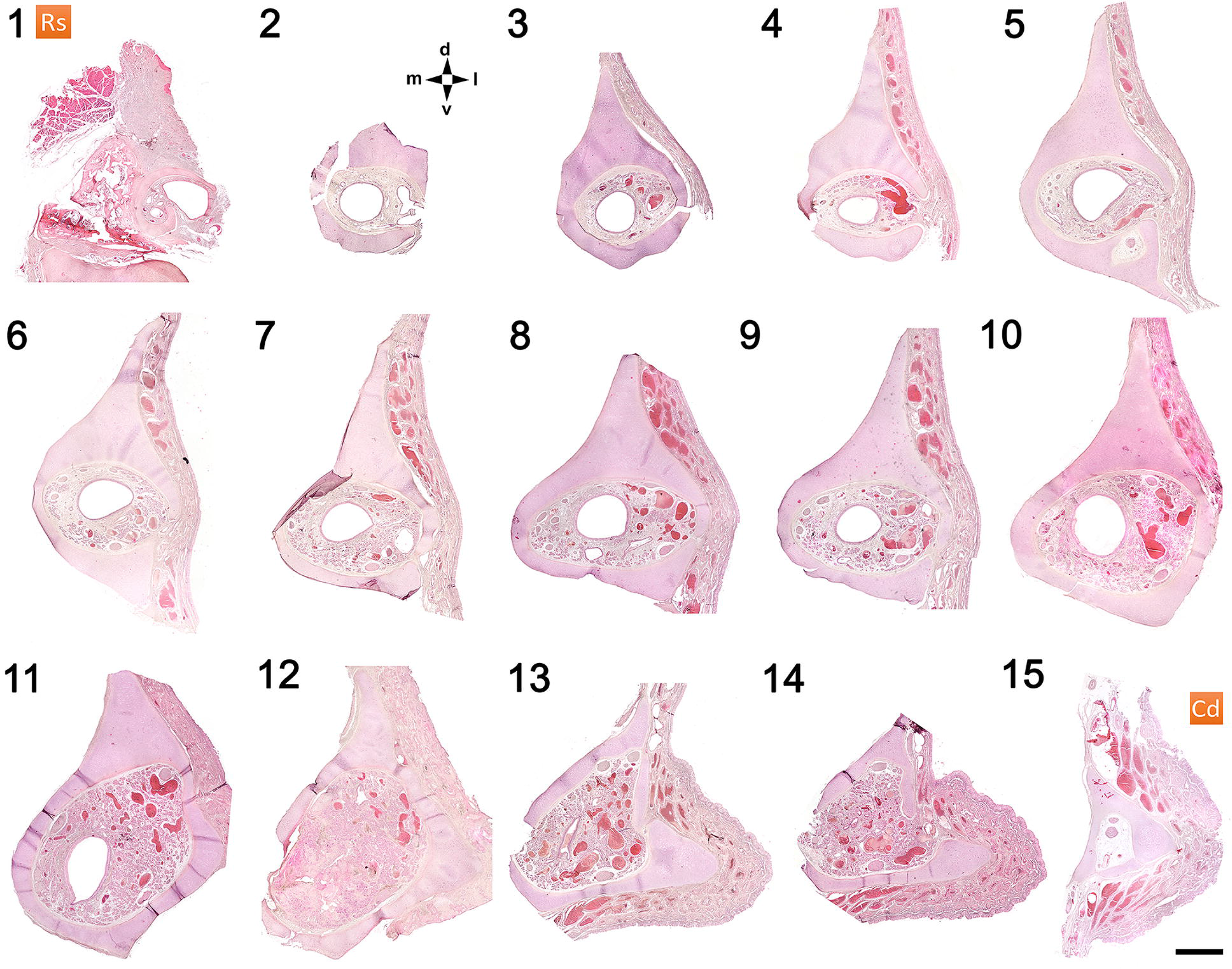
Histological transverse sections of the vomeronasal organ (VNO) at fifteen different levels numbered from rostral (1) to caudal (15) and stained by hematoxylin-eosin. Throughout the histological series, changes in the shape and dimension of the cartilage and the vomeronasal duct can be observed. Variations in the main structural features in the parenchyma of the VNO can also be noticed, especially regarding its vascular component. The first level (1) is the more rostral (Rs) studied and corresponds to the common transit of both ducts, vomeronasal and incisive, in the palatine fissure. Levels 2 and 3 show the rostral part of the VNO. The central part of the VNO corresponds to levels 5 and 6, where the two main epithelia lining of the vomeronasal duct are already differentiated. Remarkably, in the caudal part of the VNO, corresponding to level 10, the vomeronasal duct acquires a larger caliber. Level 13 is the caudal termination of the vomeronasal duct, which is no longer present in the next and last two levels. Directionality is indicated by the following: cd, caudal; d, dorsal; l, lateral; m, medial; rs, rostral; v, ventral. Scale bar: 500 μm.

The VNO cartilage is formed by a single piece of cartilage, but it varies in shape according to the level considered. Overall, it has a circular section, which is complemented by the presence along its entire length of a prominent dorsal projection which directly contacts the septal cartilage. Its rostral-most part has the clearest circular section, but from L4 to L9 the capsule cavity is slightly flattened dorsoventrally. In the caudal part of the VNO, the cartilage undergoes striking modifications. In particular, at levels L10-L12 there is a visible increase in its cavity, which coincides with the end of the vomeronasal duct. From that point, the cartilage acquires a triangular section and undergoes a rapid reduction of its lumen, such that the extent of the parenchyma progressively decreases, so that at the most caudal level considered (L15) the lumen contains only the main branches of the innervation and irrigation of the VNO. In the parenchyma, the development of the vascular component of the organ, which in the majority of sections entirely occupies its lateral region, is remarkable. This component is composed primarily of massive venous sinuses.

The serial histological study at the block corresponding to L1 (Fig. 3) allowed to determinate the existence of a functional communication between the VNO and the incisive duct (ID) (Fig. 3A). At this point the vomeronasal duct (VD) has a much narrower diameter than the ID. Both ducts increase their caliber caudally. In the parenchyma that surrounds them, there are no glands; instead, there are vast venous sinuses. Throughout the entire level L1 both ducts are enclosed by a single cartilaginous capsule whose medial branch contacts the vomer bone (Fig. 3C,E). Only in the most rostral sections the part of cartilage surrounding the vomeronasal duct is split into two distinct parts (Fig. 3A). The epithelial lining of both ducts changes remarkably throughout this rostral level. The VD is rostrally covered by squamous simple epithelium (Fig. 3A,C), but more caudally it is replaced by a pseudostratified epithelium comprising two cell layers (Fig. 3E). Two cell types are differentiated in it: large cuboidal cells and more elongated cells whose morphology recalls neuroreceptors (Fig. 3F). Rostrally, the ID consists of a well- developed stratified squamous epithelium, mainly in its lateral part (Fig. 3B), but at the level of the communication between both VD and ID is replaced by a squamous simple epithelium (Fig. 3D). At the most caudal sections of this level L1, it is replaced by a pseudostratified lining, rich in superficial caliciform cells, and similar to the respiratory mucosa (Fig. 3G).

**Figure 3.**
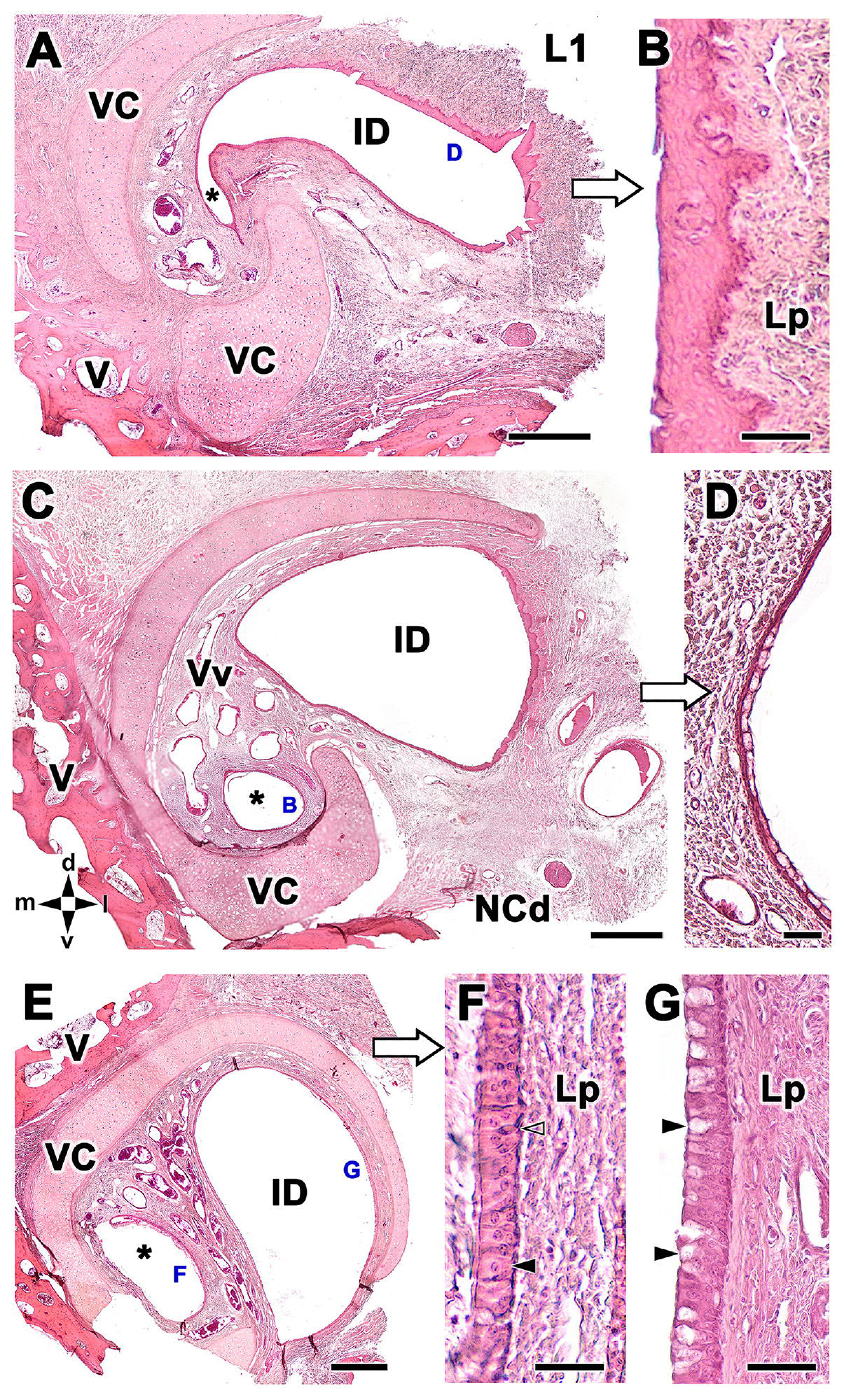
Histological study of the communication between the VNO and the incisive duct (ID). A. A rostral section in the block L1 demonstrates the point where the vomeronasal duct (*) establishes communication with the ID. At this point, the vomeronasal cartilage (VC) is split into two distinct parts. Both ducts are enclosed by a single cartilaginous capsule whose medial branch contacts the vomer bone (V). In the parenchyma, there are no glands, only large venous sinuses (Vv). The caudal nasal nerve (NCd) runs ventrally to the ID. **B.** At higher magnification, the ID is covered by a well-developed stratified squamous epithelium, mainly in its lateral part. **C.** In a more caudal section than the previous one, the vomeronasal duct is now complete, although significantly smaller than the ID. **D.** A closer look at the vomeronasal duct reveals that it is covered by squamous simple epithelium at this level **E.** A more caudal section at the L1 level shows how the vomeronasal duct has increased in size. At the same time, the nature of the lining of both the vomeronasal and incisive duct has changed, as can be seen at higher magnification in images F and G. **F.** The vomeronasal epithelium is pseudostratified and consists of two cell layers. Two cell types are differentiated: large cuboidal cells (black arrowhead) and more elongated cells (open arrowhead), which are apparently neuroreceptors. **G.** In contrast, the ID epithelium is a denser pseudostratified lining and possess superficial caliciform cells (arrowheads). Stain: hematoxylin-eosin. Lp, lamina propria. Orientation: d, dorsal; l, lateral; m, medial; v, ventral. Scale bars: A, C, E = 500 μm, B, D, F, G = 50 μm.

At both the L2 and L3 levels the VD is circular in shape (Figs. 4A, D). At L2 level laterally, the parenchyma of the organ presents large venous sinuses (Fig. 4A), whereas the rest of the parenchyma has hardly any glands and vessels, consisting mostly of connective tissue and a few small branches of the vomeronasal nerves (Fig. 4A). The lateral epithelium of the duct shows for first time basal cells and on top of them a pseudostratified epithelium formed by two layers of rounded or oval cells with nuclei densely stained by hematoxylin and individual superficial cells with translucent nuclei (Fig. 4B). The medial epithelium shows similar characteristics to those observed at the L1 level, with polyhedral cells with rounded nuclei in apical position and a simple layer of basal cells (Fig. 4C). At L3 level the vomeronasal cartilage presents a sharp dorsal projection and encircles the entire organ. In comparison with the previous level, the parenchyma presents a great development of the glandular component, that together with the venous sinuses is concentrated in the dorsolateral quadrant (Fig. 4D). The lateral epithelium is similar to that observed in the previous level (Fig. 4E), whereas there are changes in the organization of the medial vomeronasal epithelium. The previously geometrically shaped cells have been replaced by cells with elongated nuclei and cytoplasm and arranged in an incipient columnar manner. This represents a transition to the typical vomeronasal receptor epithelium (Fig. 4F).

**Figure 4.**
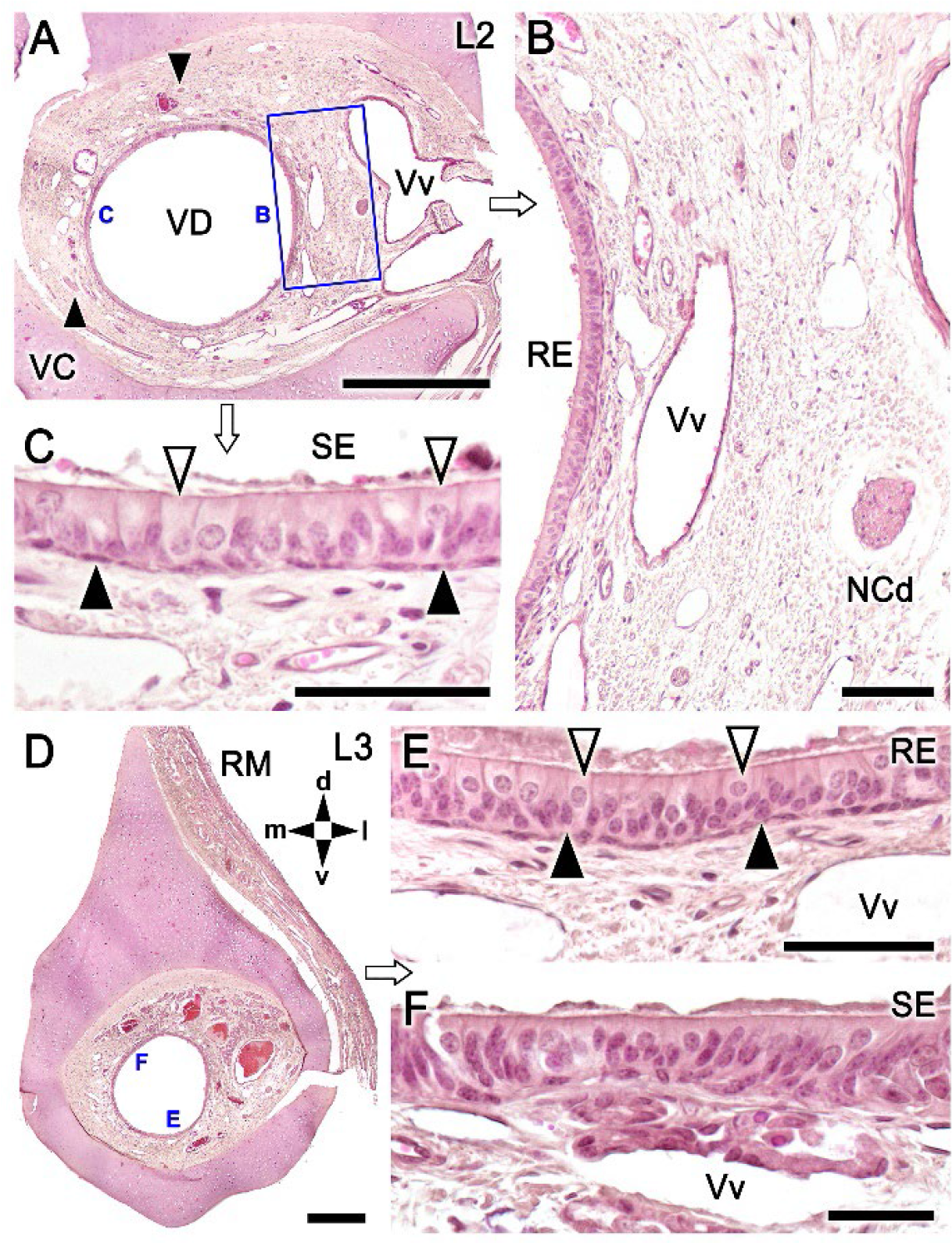
Histological study of the VNO at the rostral levels L2 (A-C) and L3 (D-F). A. At the L2 level, the VD is circular in shape. Laterally, the parenchyma of the organ presents large venous sinuses, which reach into the lateral opening of the vomeronasal cartilage. The rest of the parenchyma has hardly any glands and vessels, consisting mostly of connective tissue and a few small branches of the vomeronasal nerves (arrowheads). **B.** Enlargement of the area boxed in A, showing at higher magnification the respiratory epithelium. Basal cells are seen for the first time at this level and on top of them a pseudostratified epithelium formed by two layers of rounded or oval cells with nuclei densely stained by hematoxylin and isolated superficial cells with translucent nuclei. **C.** The medial epithelium shows similar characteristics to those observed at the L1 level. It presents polyhedral cells with rounded nuclei in apical position (open arrowhead) and a simple layer of basal cells (black arrowhead). **D.** At L3 level the vomeronasal cartilage presents a sharp dorsal projection and encircles the entire organ. In comparison with the previous level, the parenchyma presents a great development of the glandular component. Together with the venous sinuses, it is concentrated in the dorsolateral quadrant. **E.** The lateral epithelium is similar to that described in the previous level with superficial cuboidal cells (open arrowheads) and dark nucleated cells (black arrowheads) located in a basal position. **F.** There are changes in the organization of the medial vomeronasal epithelium; the previously geometrically shaped cells now have elongated nuclei and are arranged in an incipient columnar manner. This represents a transition to the typical vomeronasal receptor epithelium. RE, respiratory epithelium; RM, respiratory mucosa; SE, sensory epithelium; VD, vomeronasal duct; Vv, veins; directionality is indicated by the following: d, dorsal; l, lateral; m, medial; v, ventral. Scale bars: A, D = 500 μm, B = 100 μm, C, E, F = 50 μm.

Level L5 (Fig. 5A-E) corresponds to the central part of the VNO. As most prominent features, the vomeronasal cartilage presents a lateral opening and the presence of a ventral recess for the caudal nasal nerve (NCd) (Fig. 5A). At this point, the vomeronasal duct adopts its characteristic crescent shape, with the medial epithelium showing for first time the typical morphology of a full developed sensory epithelium. The basal cells are not very conspicuous, the somas of the neuroreceptor cells are large and rounded and have a distinct nucleus, eosinophilic cytoplasm, and a process towards the luminal surface (Fig. 5B). The sustentacular cells are more strongly stained and densely packed close to the luminal surface (Fig. 5B). In the respiratory epithelium a few polyhedral cells with translucent cytoplasm are seen in its apical zone. The medial parenchyma is devoid of glands, but contain numerous branches of the vomeronasal nerves (Fig. 5D), whereas the lateral part of the duct is connected to tubular glands that open in the dorsal commissure of the VD (Fig. 5E).

**Figure 5.**
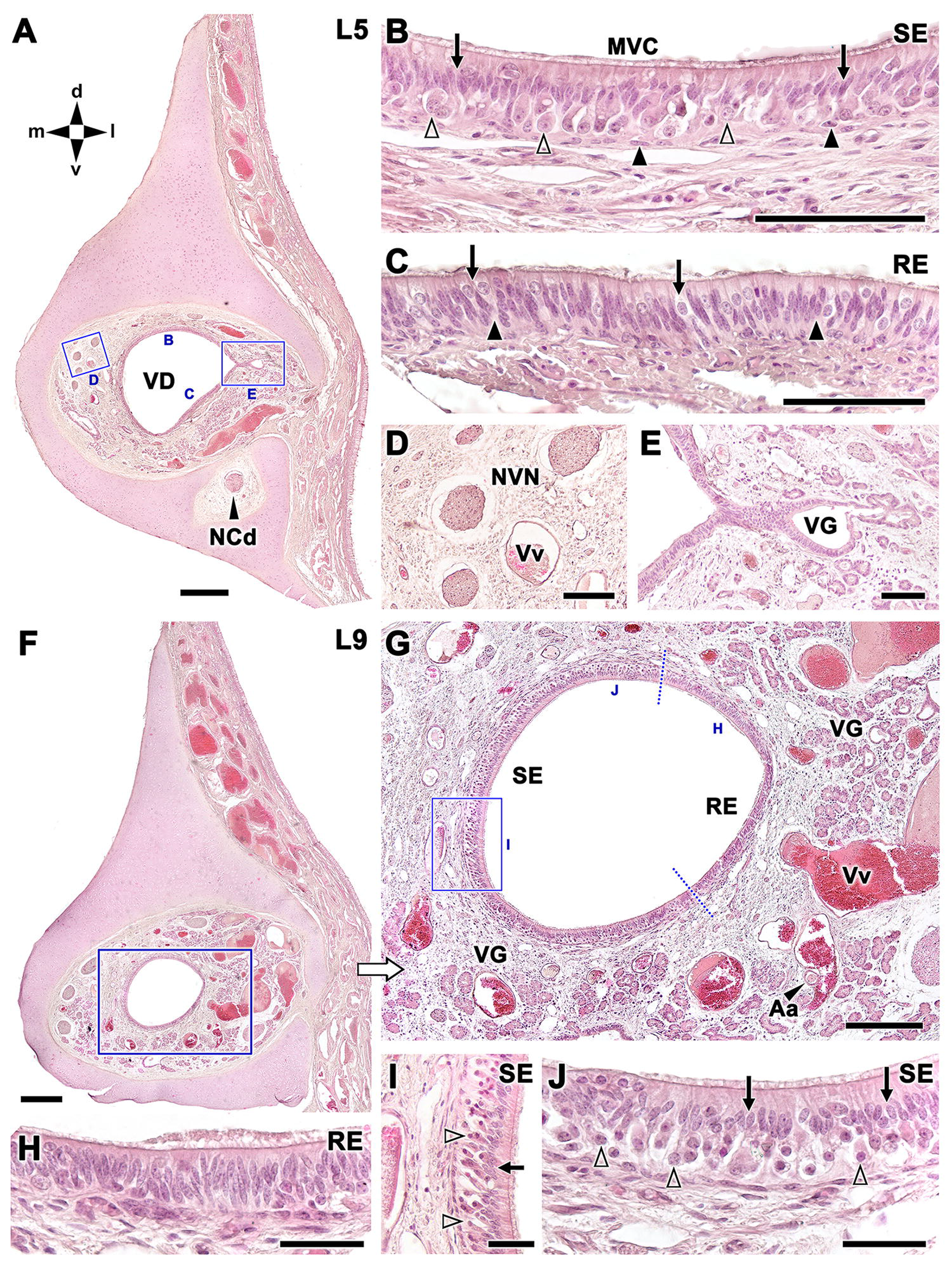
Histological features of the central (L5) and caudal (L9) parts of the VNO of the dama gazelle. A. At the central level, the more prominent features of the vomeronasal cartilage are its lateral opening and the presence of a recess for the caudal nasal nerve (NCd). At this point, the vomeronasal duct adopts its characteristic crescent shape. **B.** The morphology of the medial epithelium is typical of vomeronasal sensory epithelium. The basal cells are not very conspicuous (black arrowheads), the somas of the neuroreceptor cells (open arrowheads) are rounded and have a distinct nucleus, eosinophilic cytoplasm, and a projection towards the surface. They are large cells, in contrast to sustentacular cells, which are more heavily stained (black arrows), and packed close to the luminal surface of the epithelium. A mucomicrovillar complex (MVC) covers the luminal surface of the SE. **C.** Polyhedral cells with translucent cytoplasm are seen in the apical zone of the respiratory epithelium (arrows), whereas cells with darkly stained ellipsoidal nuclei (arrowheads) are found throughout the epithelium. **D.** The branches of the vomeronasal nerves (NVN) are highly developed and located in the medial parenchyma. **E.** The glandular tissue is located laterally in the parenchyma. It is less abundant than in the preceding level. The glands (VG) discharge their secretions into channels that open into the vomeronasal duct (VD). **F.** At a more caudal level of the VNO (L9), the VC is almost completely closed. The vomeronasal duct (VD) has reduced in diameter and its shape is again circular and shows increased development of vascular erectile tissue. The NCd is again inside the parenchyma, and numerous thick vomeronasal nerves run medially. **G.** At higher magnification (box in F), the transitions between the sensory and respiratory epithelia are marked with two dashed dotted lines. This shows that about ¾ of the total epithelial lining of the duct is of neuroreceptor nature. **H.** The respiratory epithelium (RE) is similar to that of the central level (L5), although there are fewer polyhedral cells with clear nuclei. The surface of the epithelium has numerous cilia. **I, J.** In the sensory epithelium (SE) the differentiation between neuroreceptor cells, in the basal part (open arrowheads), and sustentacular cells with dense nuclei (black arrows) is marked. On the luminal surface of the SE, the presence of microvilli is striking.). Aa, arteries; Vv, veins; directionality is indicated by the following: d, dorsal; l, lateral; m, medial; v, ventral. Scale bars: A, F = 250 μm, B = 250 μm, B-E, = 100 μm, H-J = 50 μm.

At a more caudal level, L9 (Fig. 5F-J), the VC is almost completely closed, and the VD has reduced its diameter and its shape is again circular. The parenchyma shows an increased development of vascular erectile tissue (Fig. 5F). The NCd is again inside the parenchyma, and numerous thick vomeronasal nerves run medially. Although the lumen is no longer crescent in shape, three quarters of the epithelial lining of the duct are of neuroreceptor nature (Fig. 5G). The sensory epithelium shows a clear distinction between neuroreceptor cells, in the basal part and more apical sustentacular cells with dense nuclei (Fig. 5I, J). The respiratory epithelium is similar to that of the central level (L5; Fig. 5C), although there are fewer superficial polyhedral cells.

Figure 6 summarizes the main histological features of caudal part of the VNO. At level L10 (Fig. 6A-D), the cartilaginous envelope is complete, and compared to the previous level L9 the parenchyma is remarkably enlarged due to the presence of copious glandular tissue and large blood vessels. Sensory and respiratory epithelia are clearly differentiated (Fig. 6B, C), being the sensory epithelium comparable to the previous level L9, with clearly defined neuroreceptor cells (Fig. 6D). However, it has a higher density of sustentacular cells than in L9. Level L13 (Fig. 6E-H) corresponds with the caudal extremity of the vomeronasal duct. Even at this caudalmost level the medial sensory and the lateral respiratory epithelia are clearly differentiated (Fig. 6F). In the SE, additionally to the receptor and sustentacular cells, there is also the presence of globose cells in a superficial location (Fig. 6G). The basal cells are less numerous than in previous levels.

**Figure 6.**
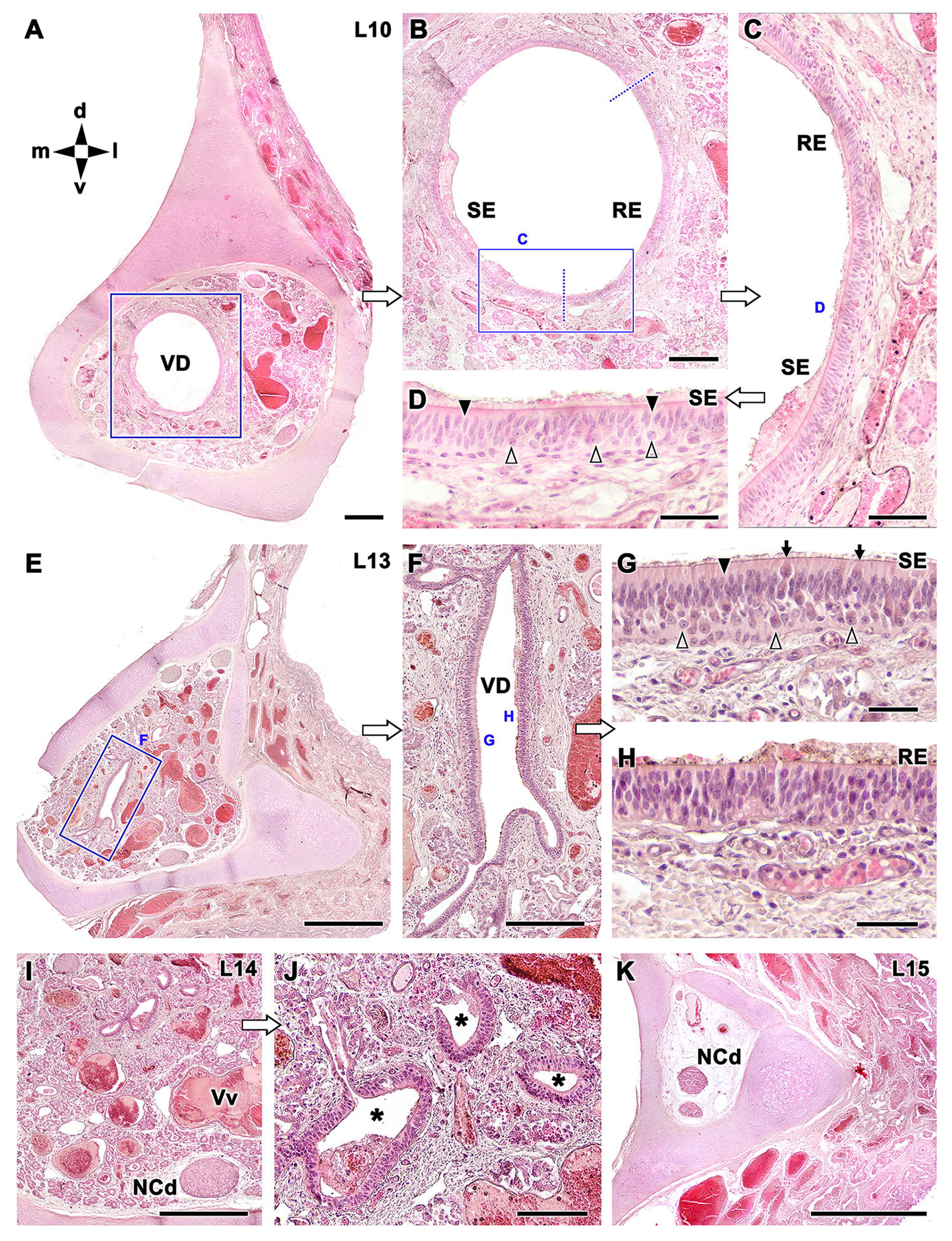
Histological features of the caudal levels of the VNO of dama gazelle. A. At level L10, the cartilaginous envelope is complete, and compared to the previous level L9 the parenchyma is remarkably enlarged due to the presence of copious glandular tissue and large blood vessels. **B, C.** Sensory and respiratory epithelia are clearly differentiated. **D.** At a higher magnification, the sensory epithelium is comparable to the previous level L9, with clearly defined neuroreceptor cells (open arrowheads). However, at this level L10 it has a higher density of sustentacular cells (black arrowheads). **E.** Level L13 corresponds with the caudal extremity of the vomeronasal duct (VD). **F.** Even at this caudalmost level the medial sensory and the lateral respiratory epithelia are still clearly differentiated. **G.** In the sensory epithelium, additionally to the receptor (open arrowhead) and sustentacular (black arrowhead), there is also the presence of globose cells in a superficial location (arrows). The basal cells are less numerous than in previous levels. **H.** The respiratory epithelium, on the other hand, maintains its structure and does not present basal cells either. **I.** At level L14 the parenchyma of the VNO is composed of venous sinuses, abundant glandular tissue, and nerve bundles, and the VD is absent. **J.** At higher magnification tubular epithelial structures (asterisk) can be detected, which correspond to the earliest epithelial invaginations that, like gloved fingers, give origin to the vomeronasal duct. These ducts are lined exclusively with respiratory epithelium. **K.** At the most caudal level studied (L15), the cartilaginous sheath is triangular and contains branches of the caudal nasal nerve (NCd), veins and an artery. RE, respiratory epithelium; SE, sensory epithelium; Vv, veins; directionality is indicated by the following: d, dorsal; l, lateral; m, medial; v, ventral. Scale bars: E, K = 500 μm; A, I = 250 μm; F = 250 μm; B, C, J = 100 μm; D, G, H = 50 μm.

The respiratory epithelium maintains its structure and does not present basal cells either (Fig. 6H). At level L14 (Fig. 6I, J) the parenchyma is composed of venous sinuses, abundant glandular tissue, and nerve bundles, and the VD is absent, but several tubular epithelial structures can be detected, which correspond to the earliest epithelial invaginations that, like gloved fingers, give origin to the VD (Fig. 6J). These ducts are lined exclusively with respiratory epithelium. At the most caudal level studied L15 (Fig. 6K), the cartilaginous sheath is triangular in shape and contains branches of the caudal nasal nerve (NCd), veins and arteries.

The study of the caudal portion of the VNO was accompanied by horizontal sections of this region (Fig. 7). This series reveals how the caudal extremity of the vomeronasal duct gives way to a large area of parenchyma rich in glandular tissue and venous sinuses flanked on both sides by the vomeronasal cartilage (Fig. 7A). Some VG are observed in the parenchyma at a ventral level, but venous sinuses predominate (Fig. 7B). At a dorsal level, the vomeronasal nerve has a remarkable number of longitudinal branches (Fig. 7C). The glandular tissue is serous in nature (Fig. 7D).

**Figure 7.**
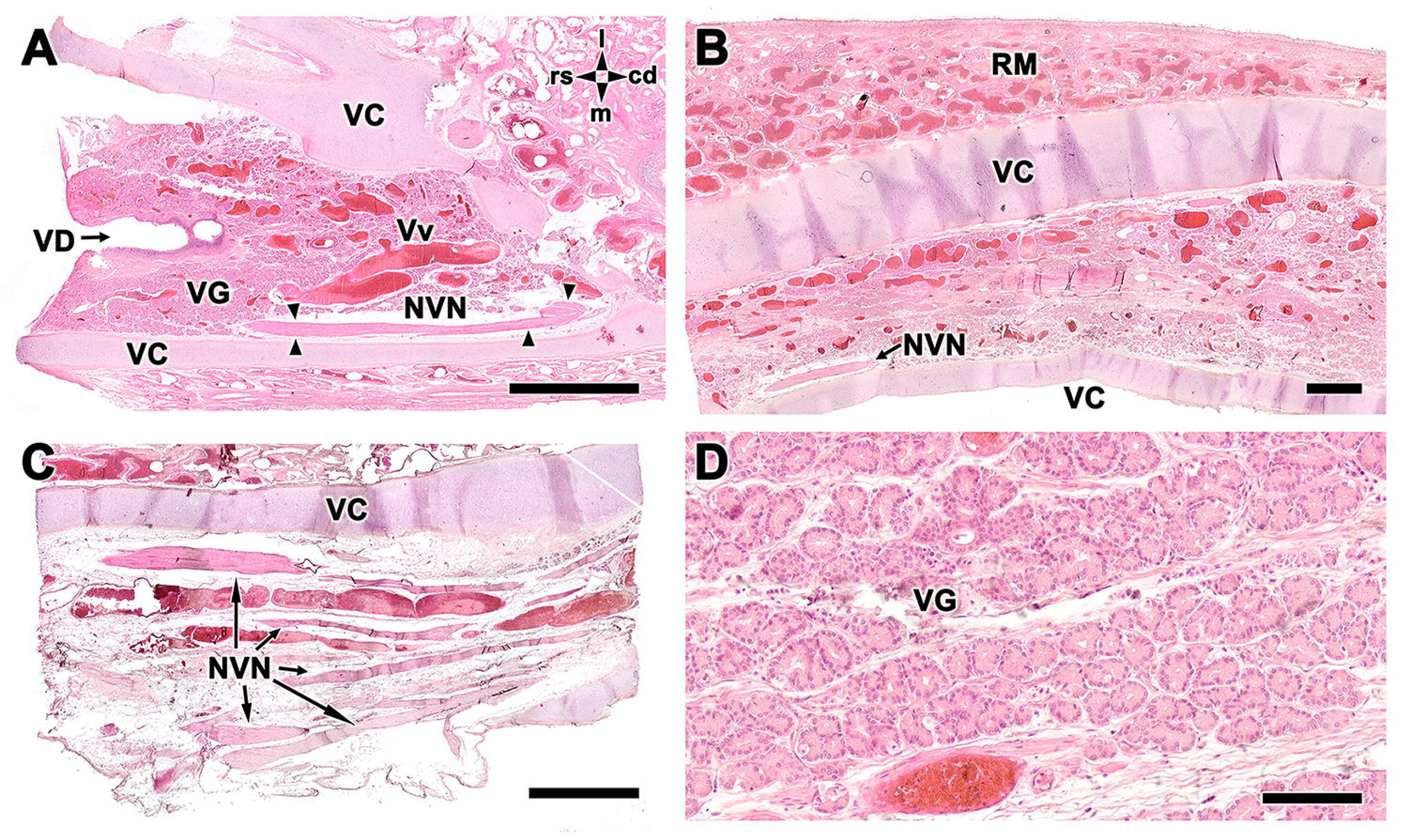
Horizontal sections of the caudal area of the VNO. A. The caudal extremity of the vomeronasal duct (VD) gives way to a large area of parenchyma rich in glandular tissue and venous sinuses (Vv) flanked by the vomeronasal cartilage for both sides. Vomeronasal nerves (NVN) are visible in the medial side parenchyma of the organ. **B.** At a ventral level, the VC remains, and some VG are observed in the parenchyma, however, venous sinuses predominate. **C.** At a dorsal level, venous sinuses are observed accompanied by numerous longitudinal branches of NVNs. **D.** Higher magnification view of the serous glandular tissue (VG) present in the caudal part of the VNO. RM, respiratory mucosa; directionality is indicated by the following: cd, caudal; l, lateral; m, medial; rs, rostral. Scale bars: A, C = 1 mm, D = 100 μm.

The serial histological study was continued with the use of specific stains for the glandular tissue, namely Alcian blue (Fig. 8A-C) and PAS (Fig. 8D-L), to identify acidic and neutral mucins in the tissue, respectively. The epithelial glands in the surface of the respiratory mucosa of the nasal cavity, the VC (Figures 8A, D, I), and the mucomicrovillar complex that covers the luminal surface of the VD epithelium were strongly stained by both stains (Fig. 8B, J). Nevertheless, there is a clear distinction between the two kinds of secretions that are produced in the glands that are located within the parenchyma of the VNO. PAS, beginning at level L3 (Fig. 8F), produced a generalized staining of the glandular component that extended all the way to the caudal end of the VNO, whereas Alcian blue only stained the central part of the VNO (level L6) and was very restricted to a small group of glands in the dorsal part of the parenchyma (Fig. 8B, C) (Fig. 8L). Therefore, focusing on the description of PAS staining, at level L3, only a small group of glands that were located ventrally were stained with PAS; however, at level L4, the VG had expanded their extension and occupied most of the dorsal parenchyma. From level L6 onwards (Fig. 8D, E), the glands encompassed the whole parenchyma surrounding the VD, filling the space among the venous sinuses. At the most caudal levels of the VD (Fig. 8H. K), where the duct is shrunk to its smallest size, PAS positive glands reached its highest level of expression.

**Figure 8.**
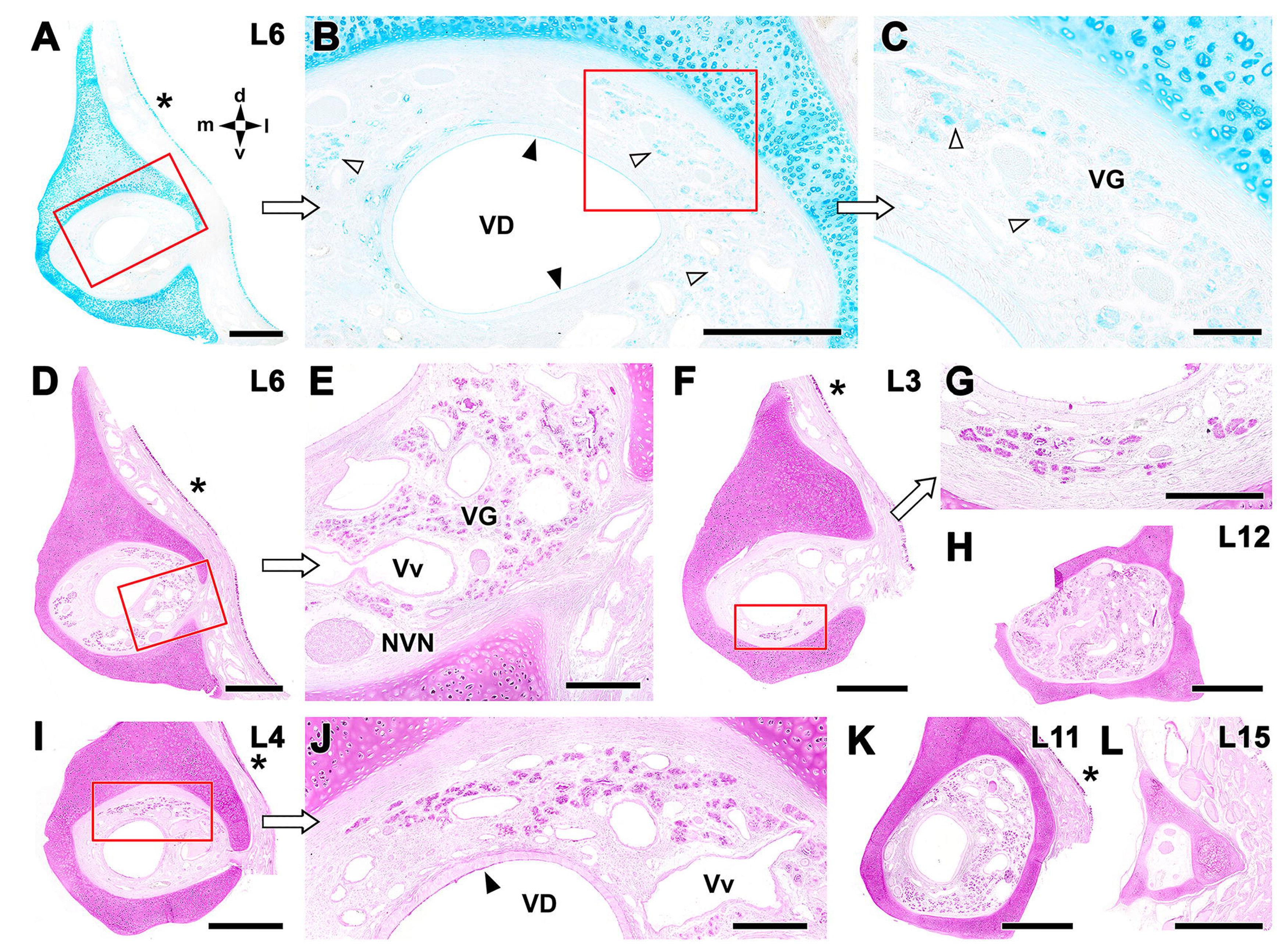
Glandular component of the gazelle dama VNO. A. Alcian blue stain marks strongly the VC and the surface of the respiratory mucosa of the nasal cavity (asterisk). **B.** Higher magnification of a section corresponding to the level L6, shows a faint staining in a small group of glands in the dorsal part of the parenchyma (open arrowheads) **C.** A magnification of the box in B shows the serous nature of the VG. The luminal surface of the RE is similarly stained (arrowhead). **D.** Contrastly, at the same level L6 PAS staining shows a high development of the glandular tissue, surrounding the entire vomeronasal duct. The VC and the mucosa of the nasal cavity (asterisk) are as well stained. **E.** The enlargement of the box in D shows the serous morphology of the VG. The veins basal lamina (Vv) and the connective tissue in the NVNs are slightly marked. **F.** The more rostral levels show a lesser development of the VG. In L3 the glands are located ventrally, and they are serous in nature **(G). I.** At level L4 they are located dorsally, occupying most of the dorsal parenchyma **(J). H, K.** At the most caudal levels of the VD (L12, L11, respectively), PAS positive glandular development reaches its maximum expression. **L**. The most caudal level of the VNO studied does not present VGs. Directionality is indicated by the following: d, dorsal; l, lateral; m, medial; v, ventral. Scale bars: A, D, F, H, I, K, L = 500 μm; E, J = 150 μm; B, C, G = 100 μm

#### 2.2 Immunohistochemical study

The immunolabeling pattern against the αi2 and αo subunits of the G proteins (Fig. 9) is especially revealing because, in the context of the VNO, the expression of these proteins is thought to correspond to the effective expression of the two major families of vomeronasal receptors, V1R and V2R, respectively.

**Figure 9.**
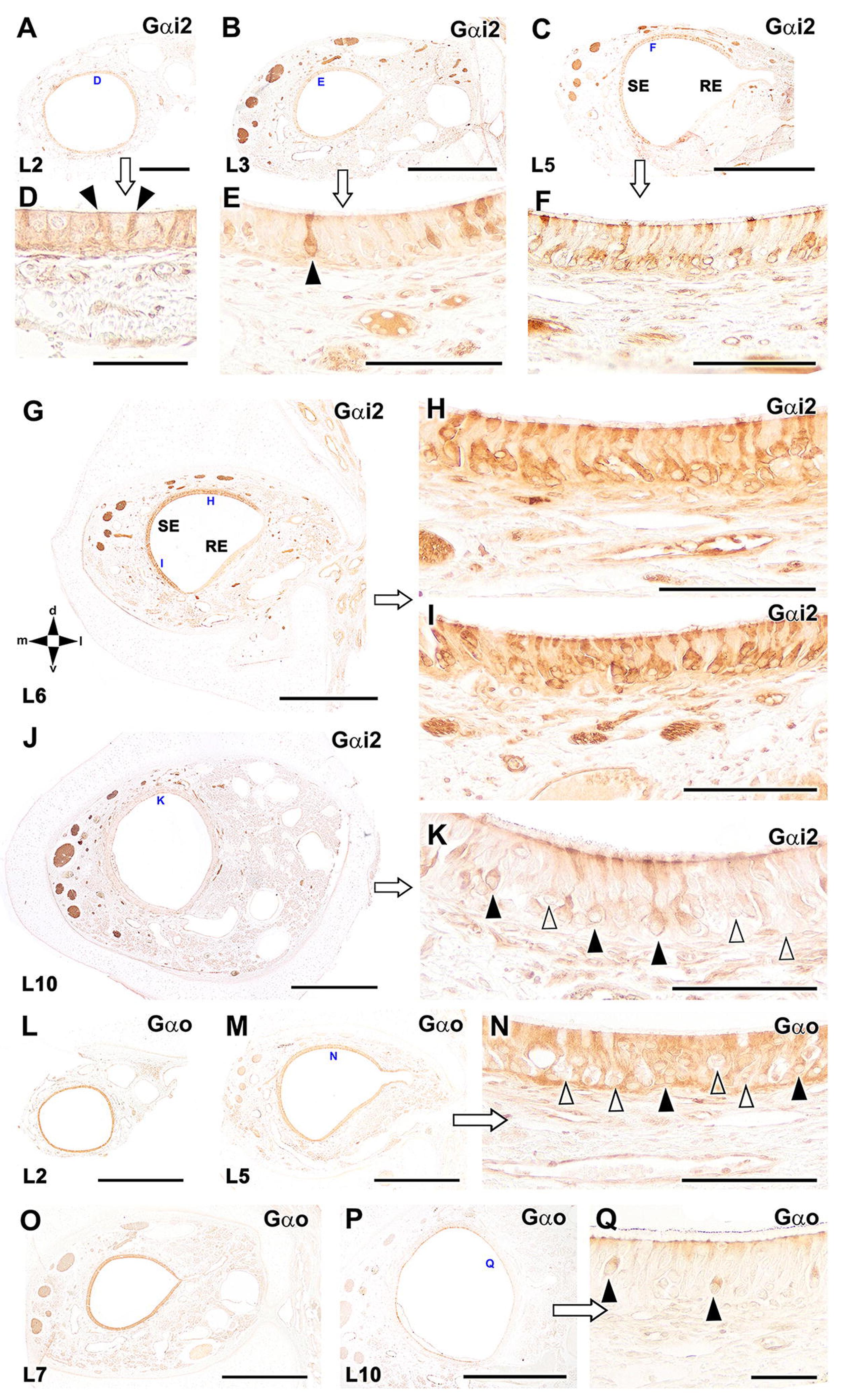
Immunohistochemical study of G-proteins in the VNO of gazelle dama. A. Gαi2 protein immunolabelling in the rostral VNO (Level L2) labels faintly the whole VD epithelium. At higher magnification **(D)** it can be appreciated how this labelling is mainly due to the presence of scattered cells with neuroepithelial morphology and strongly stained processes (arrowhead). **B.** At the next level studied (L3) the vomeronasal duct, except for its lateral part, still shows diffuse immunolabeling. This contrasts with the presence of strong immunolabeling in the vomeronasal nerves. This fact will be a constant throughout the entire VNO. At higher magnification **(E)** the epithelium still presents scattered neuroreceptor cells, but in comparison to the previous level more differentiated and with larger and strongly stained processes (arrowhead). **C.** In the central level of the VNO (L5) the difference between the immunopositive sensory epithelium and the immunonegative lateral epithelium becomes evident for the first time. At higher magnification **(F)** the sensory epithelium presents its typical features: a continuous layer of neuroreceptor-positive somata with neatly stained cytoplasm and dendritic boutons. **G.** At the L6 level where the vomeronasal duct acquires its crescent shape, the difference between the SE and the RE is more remarkable. **H, I.** the whole length of the SE presents a similar microscopic structure. **J.** The L10 level reflects an important modification at the caudal level; apparently there is hardly any positive immunolabeling along the entire perimeter of the VD epithelium. **K.** However, at higher magnification the presence of immunopositive cells (black arrowheads) scattered among immunonegative cells (open arrowheads) is observed. **L.** Immunoabelling with the anti-Gαo antibody was present in the whole VD epithelium from the more rostral level L2. **M, O**. At levels L5 and L7 the NVN are immunolabelled and there is no discrimination between lateral and medial epithelia. **N**. At a higher magnification of the SE in L5, it is apparent that not all neuroreceptor cells of the epithelium are Gαo-positive, with distinct Gαo-negative regions (open arrowheads) interspersed among immunopositive Gαo cells (black arrowheads). **P.** In level L10, there is no immunostaining of the VD epithelium, as it was the case with Gαi2. **Q.** However, higher magnification reveals the presence of Gαo-positive neuroreceptor cells, but only in the lateral RE (arrowheads). Directionality is indicated by the following: d, dorsal; l, lateral; m, medial; v, ventral. Scale bars: C, G, J, L, M, O, P = 500 μm; A, B = 250 μm; E, F, H, I, K, N = 100 μm; Q = 50 μm; D = 25 μm

Gαi2 protein immunolabelling (Fig. 9 A-K) varies markedly throughout the VNO (Level L2). At the most rostral level L2 it labels faintly the whole VD epithelium (Fig. 9A). This labelling is mainly due to the presence of scattered cells with neuroepithelial morphology and strongly stained processes (Fig. 9D). At the next level studied (L3) appears a strong immunolabeling in the vomeronasal nerves (Fig. 9B), which will be a constant throughout the entire VNO. At this L3 level the neuroreceptor cells are still scattered, but they possess large and strongly stained processes (Fig. 9E). In the central levels of the VNO (L5, L6) the difference between the immunopositive sensory epithelium and the immunonegative lateral epithelium becomes evident (Figs. 9 C, G). In both cases the SE presents its typical features: a continuous layer of neuroreceptor- positive somata with neatly stained cytoplasm and dendritic boutons (Figs. 9 F, H, I). The L10 level reflects an important modification as apparently there is hardly any positive immunolabeling along the entire perimeter of the VD epithelium (Fig. 9J). However, at higher magnification the presence of Gαi2 immunopositive cells scattered among Gαi2 immunonegative cells is observed (Fig. 9K).

The immunolabelling against Gαo (Fig. 9L-Q) was present in the whole VD epithelium from the more rostral level L2 (Fig. 9L). At the central levels L5 and L7 the NVN are immunolabelled and there is no discrimination between lateral and medial epithelia (Fig. 9M, O). Not all neuroreceptor cells of the epithelium are Gαo-positive, as immunopositive Gαo cells are interspersed among identifiable Gαo-negative regions (Fig. 9N). At the more caudal level L10, there is no immunostaining of the VD epithelium, as it was the case with Gαi2 (Fig. 9P). However, higher magnification reveals the presence of Gαo-positive neuroreceptor cells, but only in the lateral RE (Fig. 9Q).

Neuron-specific enolase (NE), a neuronal-specific maturity marker, produces a distinctive label in both the vomeronasal nerves (NVN) and the sensory epithelium (SE), but its distribution varies depending on the level of study (Fig. 10 A-G). In the rostral part of the VNO (Level L3) there are already mature neuroreceptor cells in the dorsomedial SE (Fig. 10A). The neuroreceptor cells are interspersed throughout the SE (Fig. 10D). At central levels L5 (Fig. 10B) and L6 (Fig. 10E) there is a clear differentiation between the immunopositive medial SE and the immunonegative lateral RE. Immunolabeling is present on both the NVNs and the NCd nerve, but the latter is less intense. In the SE, neuronal somas of varying sizes and degrees of immunolabeling spread over the entire basal layer (Fig. 10C), occasionally appearing neurons whose somas are atypically located next to the lumen (Fig. 10H). At the caudal level L10 the density of the immunolabelling in the SE has decreased (Fig. 10F), but higher magnification of the SE (Fig. 10G) shows a high density of NE-immunopositive neuroreceptor cells. The OMP antibody (Fig. 10I-N) produces diffuse immunolabelling in the VD epithelium throughout the rostral (L2, L3) and central levels (L6) of the VNO (K). At higher magnification it is noticeable that immunolabelling is more intense in individual neuroreceptor cells (Fig. 10L-N). Anti-OMP does not produce immunolabelling in the VD epithelium from the caudal level L10.

**Figure 10.**
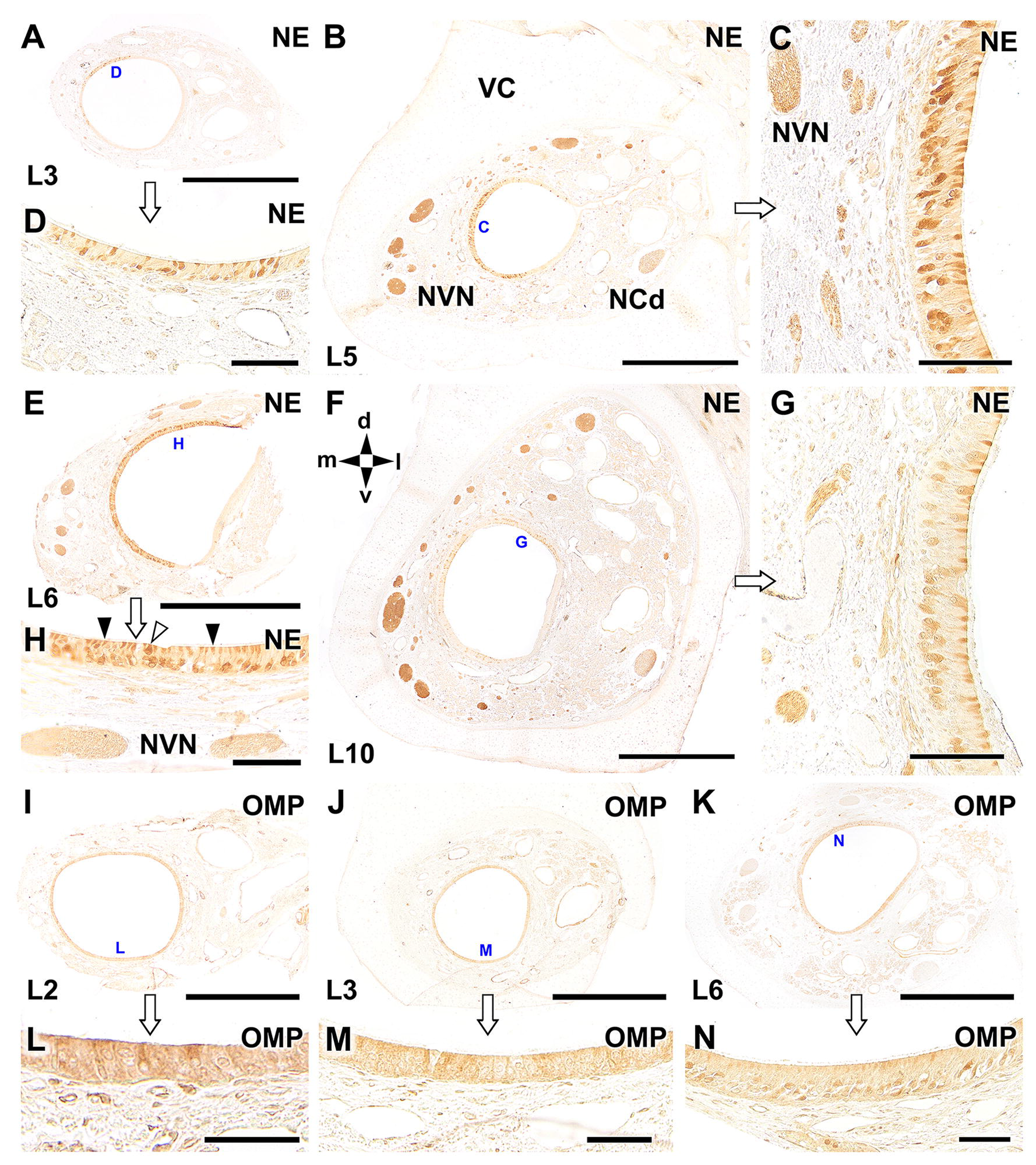
Analysis of neuronal maturity in the VNO of the dama gazelle. Neuron-specific enolase (NE), a neuronal maturity marker, produces a distinctive label in both the vomeronasal nerves (NVN) and the sensory epithelium (SE), but its distribution varies depending on the level of study. **A.** In the rostral part of the VNO (L3) there are already mature neuroreceptor cells in the dorsomedial SE. At higher magnification **(D)** the neuroreceptor cells are shown to be interspersed throughout the SE. **B.** At the central level L5 there is a clear differentiation between the immunopositive medial SE and the immunonegative lateral RE. Immunolabeling is present on both the NVNs and the NCd nerve, but the latter is less intense. Higher magnification of the SE **(C)** shows how neuronal somas of varying sizes and degrees of immunolabeling spread over the entire basal layer. **E.** The next caudal level, L6, shows a similar organization of the SE. Higher magnification of the SE **(H)** shows how the apical zone of the SE contains the dendritic knobs of the receptor cells (black arrowheads) and individual neurons whose somas are atypically located next to the lumen (open arrowhead). **F.** At the caudal level L10 the density of the immunolabelling in the SE has decreased, but higher magnification of the SE **(G)** shows a high density of NE-immunopositive neuroreceptor cells. **I, J.** The OMP antibody produces diffuse immunolabelling in the VD epithelium throughout the rostral (L2, L3) and central levels (L6) of the VNO **(K)**. **L, M, N.** At higher magnification in these levels three it is noticeable that immunolabelling is more intense in individual neuroreceptor cells. Directionality is indicated by the following: d, dorsal; l, lateral; m, medial; v, ventral. Scale bars: A, B, E, F, I-K = 500 μm; C, D, G, H, L-N = 100 μm.

The calcium-binding protein PGP 9.5 allows characterization of the neuroreceptor elements of the VNO (Fig. 11A-E). At a central level (L7) anti-PGP 9.5 immunolabels the NVN and the SE (Fig. 11A). The immunolabelling comprises all the neuroreceptor cells in the basal layer and their dendritic knobs (Fig. 11B). At the more caudal level L11, the general pattern of immunolabelling is similar (Fig. 11C), but compared to the level L7, the immunopositive cells in the SE are not so uniformly arranged, even sometimes they are aggregated in clusters (Fig. 11D). In the RE, anti-PGP 9.5 only labels a few isolated cells (Fig. 11E). Anti-cytokeratin 8 (CYK8) is a general marker for epithelial cells (Fig. 11 F-J). It produces a strong, almost homogeneous staining in the SE and RE epithelia all through the VNO, from its more rostral levels L2 (Fig. 11F) and L3 (Fig. 11G). However, some few isolated cells are not immunolabelled (Fig. 11H, J).

**Figure 11.**
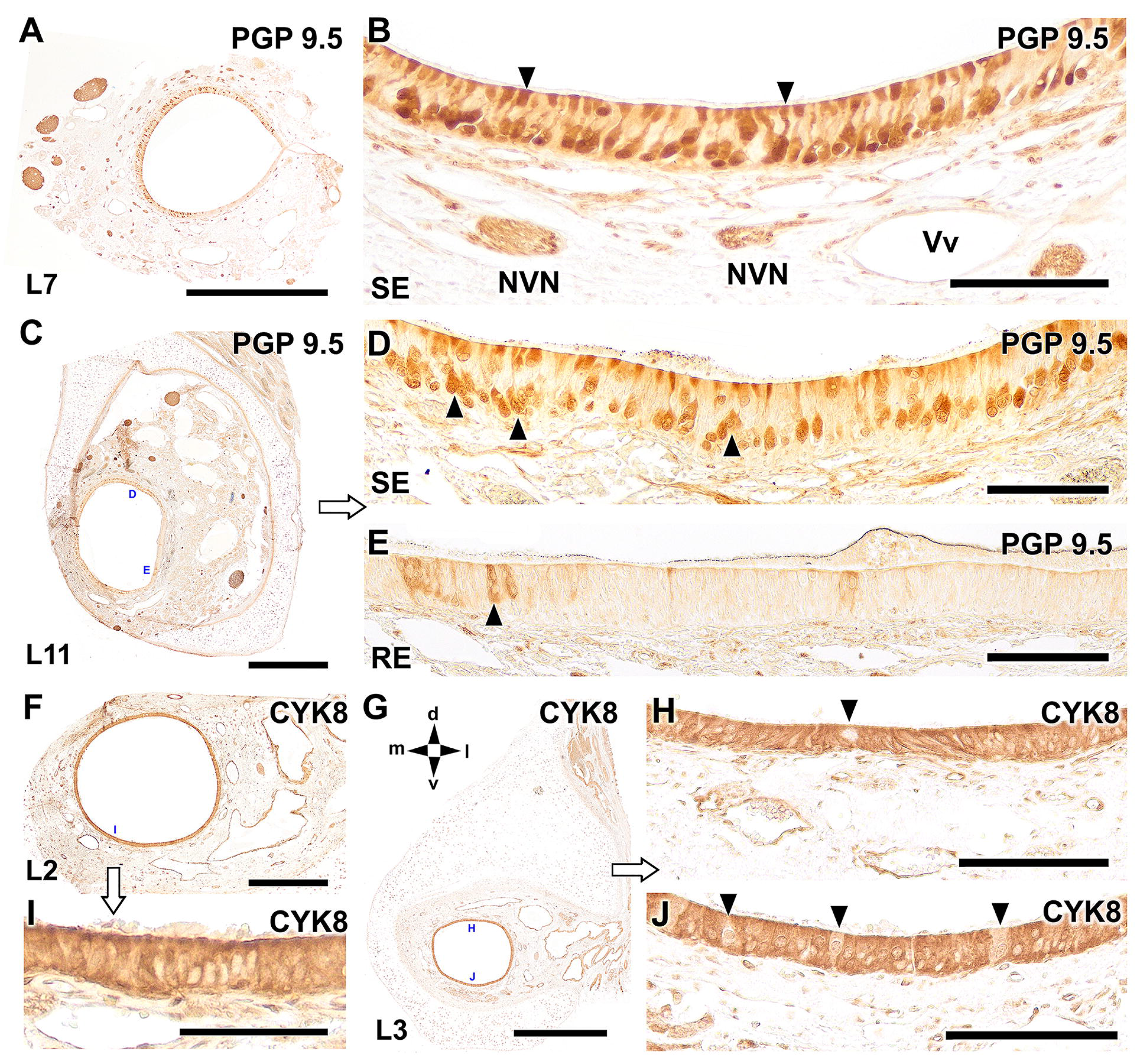
Characterization of the vomeronasal epithelium of the dama gazelle with anti-PGP 9.5 and CYK8. A. The calcium-binding protein PGP 9.5 allows characterization of the neuroreceptor elements of the sensory epithelium (SE). At a central level L7 anti-PGP 9.5 labels the vomeronasal nerves (NVN) and the SE. **B.** The immunolabelling comprises all the neuroreceptor cells in the basal layer and their dendritic knobs (arrowheads). **C.** At the more caudal level L11, the general pattern of immunolabelling is similar. **D.** There is a remarkable presence of PGP 9.5 immunopositive cells in the SE, but compared to the previous level depicted, the immunopositive cells are not so uniformly arranged, even sometimes they are aggregated in clusters (arrowheads). **E.** In the RE, anti-PGP 9.5 only labels a few isolated cells (arrowhead). **F.** Anti-cytokeratin 8 (CYK8), a general marker for epithelial cells, produces a strong, almost homogeneous staining in the SE all through the VNO, even in its more rostral levels L2 **(F)** and L3 **(G)**. **H, I, J.** In the VD epithelium, there are a few isolated cells that are not immunolabelled (arrowheads). Directionality is indicated by the following: d, dorsal; l, lateral; m, medial; v, ventral. Scale bars: A, C, G = 500 μm; F = 250 μm; B, D, E, H, J = 150 μm; I = 50 μm.

Anti-calbindin (Fig. 12A, B) stains the NVN and the SE. B. Most of the neuroreceptor cells in the basal SE are immunopositive (Fig. 12A), but in the apical layers many areas remain unstained (Fig. 12B). Anti-calretinin (Fig. 12C-E) produces a similar pattern of immunolabelling to CB, strongly labelling the NVN and the SE (Fig. 12 C, D). Only a few neuroreceptor cells remain unstained (Fig. 12E). Anti-secretagogin (Fig. 12F, G, H, J, L) produces labelling in both vomeronasal epithelia and mainly in a basal localization. In contrast to the previous markers, anti-SG does not mark projections or dendritic boutons and produces a more diffuse pattern (Fig. 12H, L). In the caudal level L10 anti-SG does not stain the VD epithelia (Fig. 12J). Anti-parvalbumin (Fig. 13I, K) marks diffusely the NVNs and vomeronasal epithelia all through the VNO.

**Figure 12.**
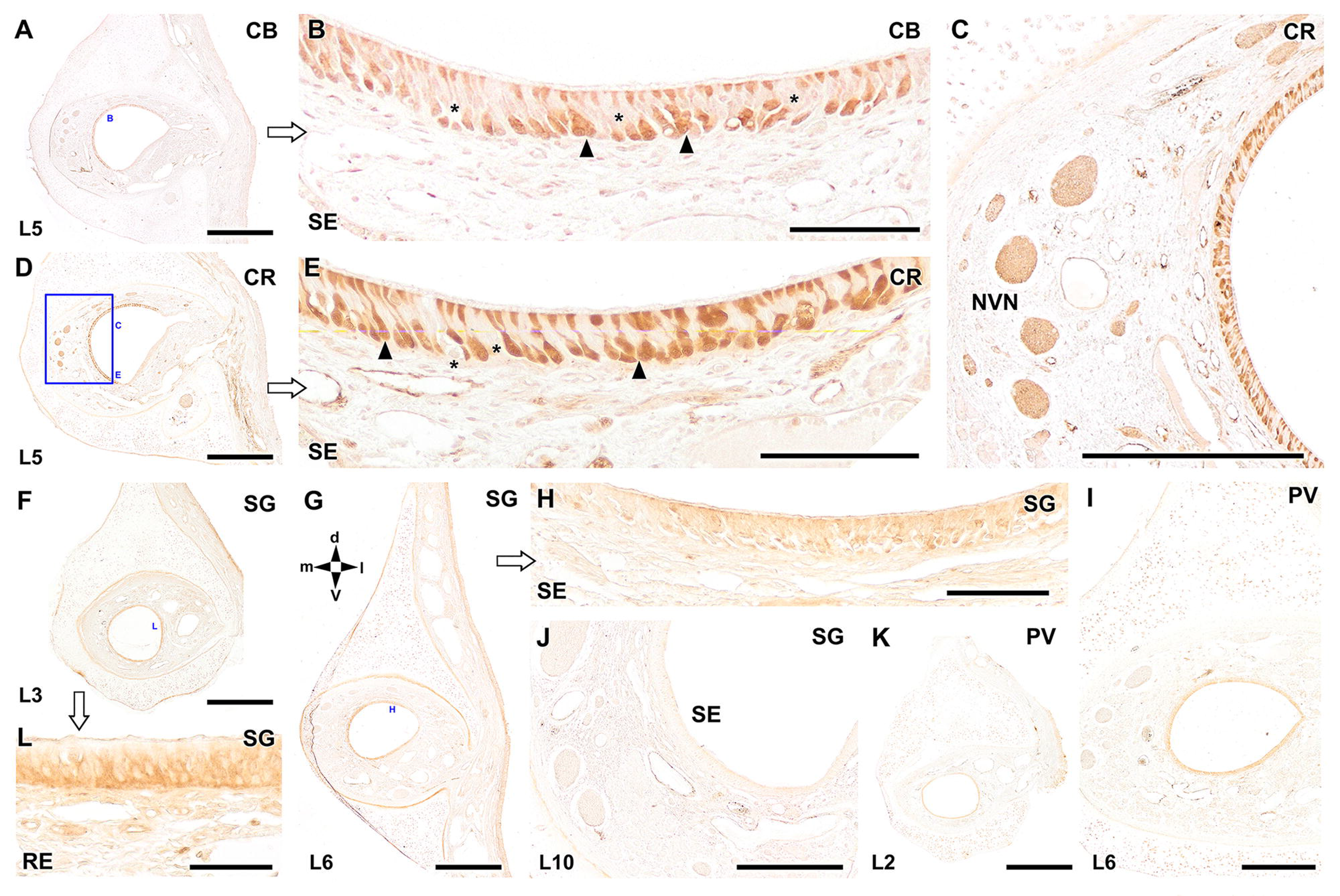
Characterization of the dama gazelle VNO by calcium-binding proteins. A. Anti-calbindin (CB) stains the vomeronasal nerves and the SE. **B.** Most of the neuroreceptor cells in the basal SE are immunopositive (arrowheads), but in the more apical layers many areas remain unstained (asterisks). **C, D.** Anti-calretinin (CR) produces a similar pattern of labelling to CB, strongly labelling the NVNs and the SE. **E.** Only a few neuroreceptor cells remain unstained (asterisk). **F, G.** Anti-secretagogin (SG) produces labelling in both vomeronasal epithelia and mainly in a basal localization **(H, L)**. In contrast to the previous markers, SG does not mark projections or dendritic boutons and produces a more diffuse pattern. In the caudal level L10 anti-SG does not stain the VD epithelia. **I, K.** Anti-parvalbumin (PV) also diffusely marks NVNs and vomeronasal epithelia. Directionality is indicated by the following: d, dorsal; l, lateral; m, medial; v, ventral. Scale bars: A, D, E, F, G, K = 500 μm; = C, I, J = 250 μm; B, E, H = 100 μm; L = 50 μm

**Figure 13.**
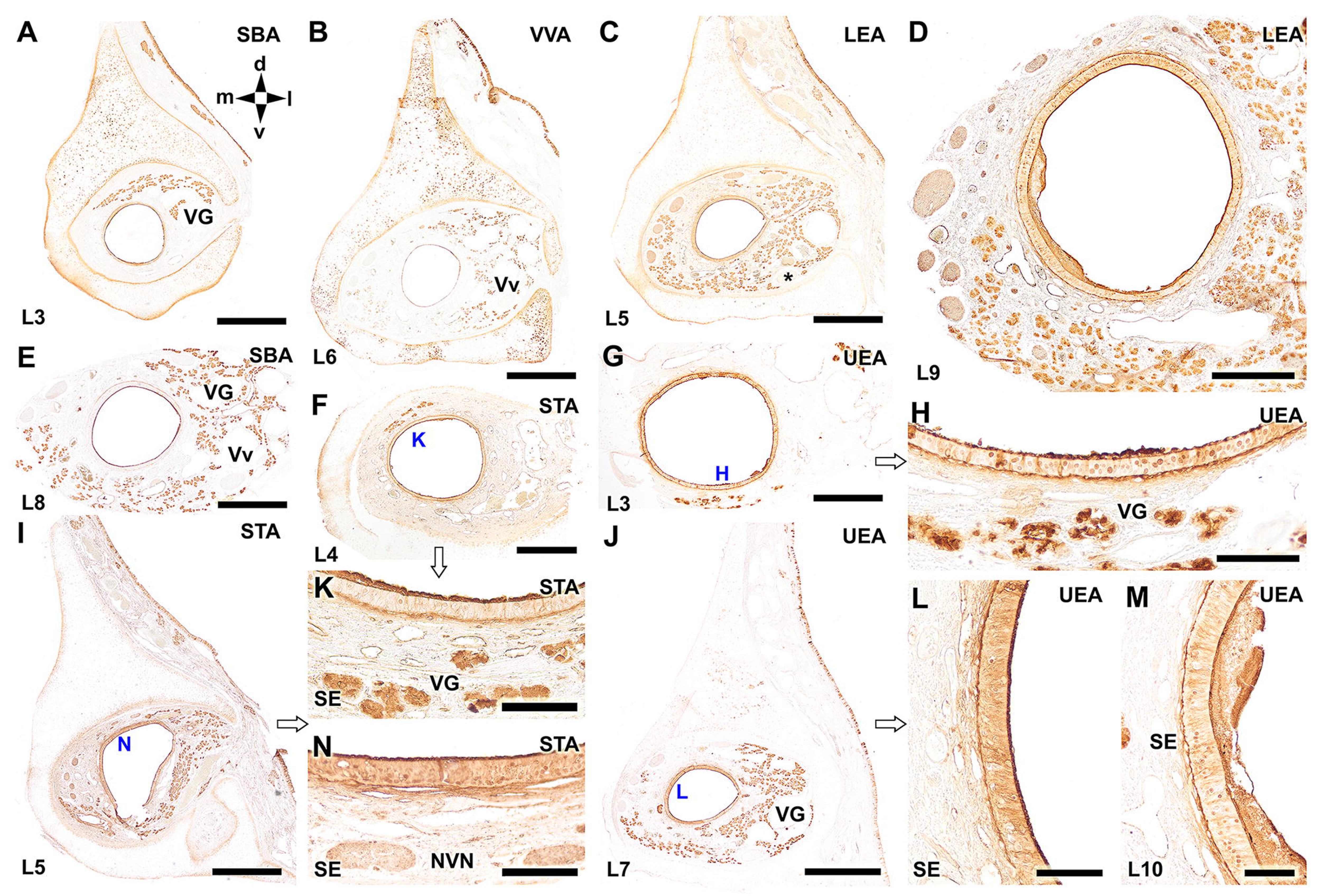
Lectin-histochemical study of the dama gazelle VNO. All the lectins employed labelled the vomeronasal glands (VG) and the muco-microvillar complex, i.e. the superficial border of both vomeronasal epithelia. **A, E**. Labelling of the VNO at levels L3 and L8, respectively with the lectin SBA (*Glycine max*). In both cases the VG are strongly labelled. **B.** VVA (*Vicia villosa*) labelling at a central level L6 of the VNO shows a faint labelling of the VG. **C, D.** LEA (*Lycopersicon esculentum*) labels the vomeronasal epithelia, the NVNs, and the VG. LEA does not stain the caudal nasal nerve (asterisk in C). **F, I, K, N.** STA (*Solanum tuberosum*) lectin labelling at central levels (L4, L5) of the VNO stains the vomeronasal epithelia, vomeronasal nerves (NVN) and VG. **G, H, J, L, M.** UEA (*Ulex europaeus*) lectin stains vomeronasal epithelia, VG, but hardly stains NVNs. Vv, veins; directionality is indicated by the following: d, dorsal; l, lateral; m, medial; v, ventral. Scale bars: A-C, E, I, J = 500 μm; D, F, G = 250 μm; H = 100 μm; K-N = 50 μm.

#### 2.3. Lectin histochemical study

All the lectins employed labelled the vomeronasal glands (VG) and the muco- microvillar complex, i.e. the superficial border of both vomeronasal epithelia (Fig. 13A- M). Labelling of the VNO at levels L3 and L8 with the lectin SBA (*Glycine max*), strongly labelled the VG (Fig. 13A, E). VVA (*Vicia villosa*) labelling at a central level (L6) of the VNO produced a faint labelling of the VG (Fig. 13 B). LEA (*Lycopersicon esculentum*) labeled the vomeronasal epithelia, the NVNs, and the VG. LEA did not stain the caudal nasal nerve (Fig. 13 C, D). STA (*Solanum tuberosum*) lectin labelling at central levels (L4, L5) of the VNO (Fig. 13F, I) stained the vomeronasal epithelia, vomeronasal nerves (NVN) and VG (Fig. 13 K, N). UEA (*Ulex europaeus*) lectin stained both the vomeronasal epithelia and VG (Fig. 13 G, H, J, L, M), but did not stain the NVNs at any level (Fig. 13 G, J).

The results of the immunohistochemical study are schematically summarized in Table 2.

**Table 2.**
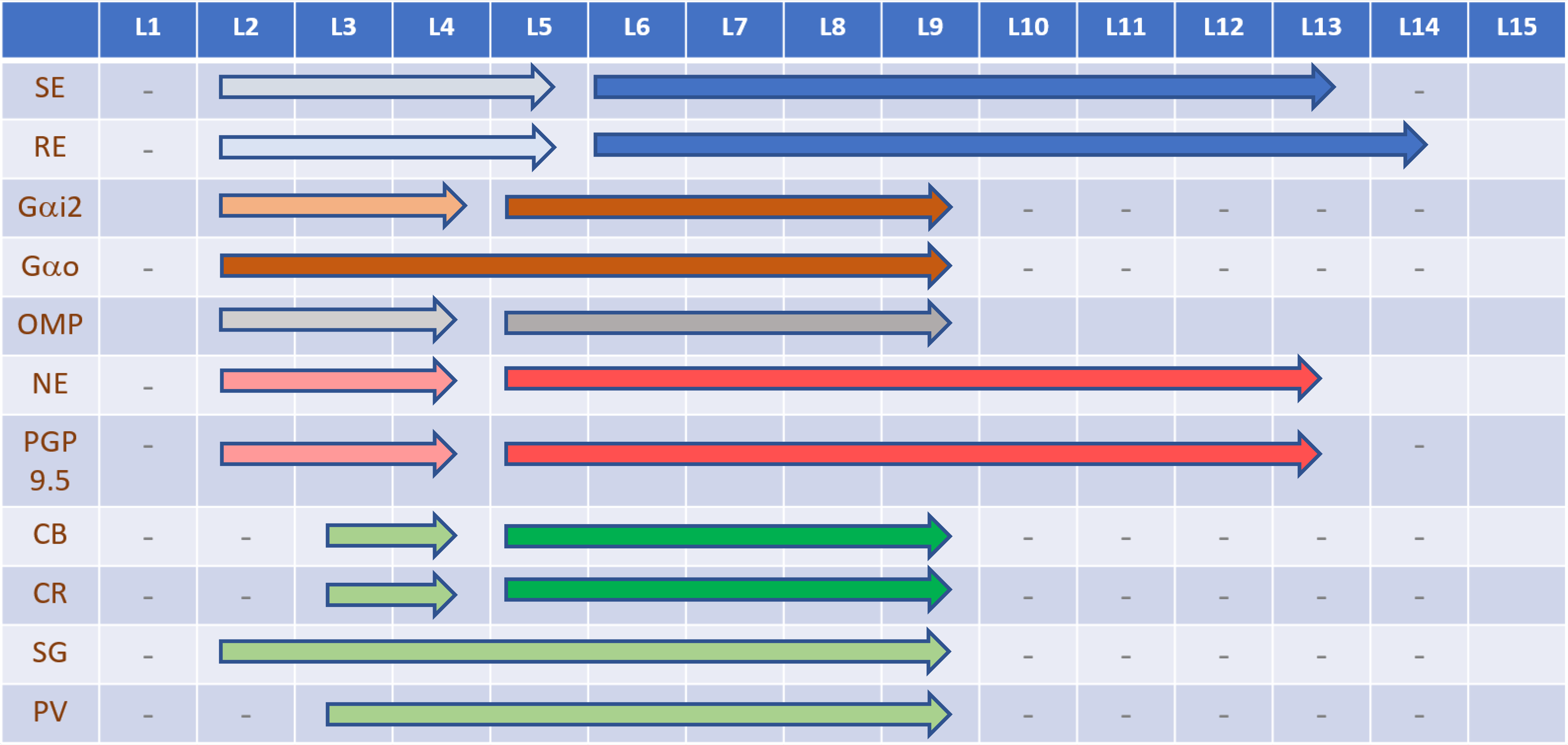
Schematic representation of neuromarker expression in the vomeronasal sensory epithelium along the rostrocaudal axis of the VNO. The occurrence of sensory epithelium and respiratory epithelium along the VD is shown in blue. The presence of a fully differentiated epithelium is shown in dark blue, while light blue shows the presence of an epithelium that has not yet adopted the typical characteristics of both sensory and respiratory epithelium. Brown shows the expression of the two subunits of the G proteins, Gαo and Gαi2 in the SE, with dark brown corresponding to full expression in the epithelium and light brown to the diffuse presence of labeled neuroreceptor cells. Dark and light gray represent the OMP expression pattern. Calciumbinding proteins are represented in dark green, in those levels where the epithelium shows a strong immunolabeling and in light green where their expression is lower

## 4. DISCUSSION

Habitat loss, overhunting, competition with livestock and climatic change have seriously compromised the survival of the gazelle dama. Maintaining a healthy and sustainable population of endangered species like this helps to preserve the overall biodiversity and ecological balance of an ecosystem. For this purpose, it is crucial to improve the welfare of gazelle dama kept in captivity. The chemical communication established between individuals of the same species can play a significant role in their reproductive behavior. This communication can involve pheromones, which are chemical substances released by one individual that affect the behavior or physiology of another individual of the same species (Liberles, 2014). In many mammals, pheromones play a crucial role in mate attraction, selection, and mating behavior. For example, female mice release pheromones that attract male mice and increase their sexual behavior (Fu et al., 2015), while male hamsters release pheromones that stimulate female ovulation and receptivity (Richardson et al., 2004). Although nowadays there is plenty of scope for the application of cattle pheromones (Rekwot et al., 2001; Landaeta-Hernández et al., 2023) in the case of the management of the reproduction of dama gazelle it requires the previous neuroanatomical and morphofunctional characterization of its vomeronasal organ, which has been approached for first time, to our knowledge in this work. Given the wide and recognized morpho-functional diversity of the mammalian vomeronasal system (Wysocki, 1979; Meisami & Bhatnagar, 1998; Smith et al., 2001; Smith, Garrett, et al., 2011), the following discussion will contextualize the results obtained on the interspecific variability of the VNO of ruminants and mammals in general. This will allow us to obtain further insight into the ecology, evolution, and conservation biology of this species.

Prior studies of the VNO suffer in many cases from being circumscribed to the central part of the organ, ignoring the changes it undergoes along its rostrocaudal axis. In contrast, there are a significant number of studies that evaluate the spatial modifications by performing serial macroscopic and histological sections that allow characterizing the most striking anatomical and histological changes (Bertmar, 1981; Salazar et al., 1995; Salazar, Lombardero, Cifuentes, et al., 2003; Vedin et al., 2010; Torres et al., 2020; Ortiz-Leal et al., 2020; Kondoh et al., 2021; Tomiyasu et al., 2022). In the case of the dama gazelle, we have extended this approach including the spatial study of the immunohistochemical and lectin-histochemical features of the vomeronasal duct and nerves. This has allowed us to integrate the morphological and functional observations along the entire VNO.

### 4.1 Macroscopic anatomy

The external morphology of the vomeronasal organ is defined by the shape of its external capsule. In the dama gazelle it is of a cartilaginous nature, encompasses the entire VNO and is topographically related in its medial wall with the vomer bone, features common to all the ruminants studied. Its shape is relatively peculiar since the most typical conformation in ruminants is oval, the major axis being dorso-ventral (Jacobs et al., 1981; Hart et al., 1988; Vedin et al., 2010; Park et al., 2014a; Ibrahim et al., 2013), whereas in the case of the dama gazelle the predominant axis is mostly lateromedial. This does not preclude notable variations along the rostro-caudal axis of the organ. When these variations are compared with the ruminant species studied in complete histological series, cow and sheep, it is observed that of the three species, the dama gazelle is the one that presents a greater morphological variation, superior to that observed in the cow (Salazar et al., 2008) and even more if it is compared with the VNO of the sheep, very homogeneous in all its length (Salazar et al., 2007). Another remarkable feature is the presence of indentations in the lateral wall of the cartilage for the passage of vessels and nerves. The three species present a wide lateral communication in its most rostral levels, while in its central part only the cow presents a wide lateral gap and the sheep lacks it (Salazar et al., 2007, 2008). In the case of the dama gazelle, this gap is moderately present, although it coincides with the formation of an internal recess that exclusively accommodates the main trunk of the caudal nasal nerve.

Less data exists on how the vomeronasal duct communicates with the external environment because determining this with accuracy requires the completion of histological series of decalcified samples. Our observation of the dama gazelle is consistent with the evidence available for other ruminants, both bovine, such as cow and sheep (Salazar et al., 2007, 2008), and non-bovine, such as the moose (Vedin et al., 2010). In all cases the vomeronasal duct opens into the duct of the incisive duct, to which it is incorporated on its medial side, both sharing the same cartilaginous capsule. Regarding the incisive papilla, we have found the presence of a functional papilla, which is of special relevance, because remarkably this is not a feature common to all Antilopinae, lacking of an incisive papilla the three species of alcelaphine antelopes studied by Hart et al. (1988).

### 4.2 Microscopic anatomy

#### 4.2.1 Vomeronasal epithelia

The extensive serial histological study carried out has allowed us to determine the existing differences along the rostro-caudal axis of the organ in terms of the epithelial lining of the vomeronasal duct, a component of special importance because it is in its medial epithelium where the neuroreceptor cells responsible for the detection of semiochemicals are located (Dennis et al., 2004; Mombaerts, 2004) and in its lateral respiratory epithelium where are located the cells producing the mucus necessary for the functioning of chemoreception (Breipohl et al., 1979; Takami et al., 1995) and growth factors modulating the neuroreceptor epithelium itself (Getchell et al., 2002)

The SE in its rostral junction with the incisive duct is covered by squamous simple epithelium which is soon replaced by a pseudostratified epithelium comprising two cell layers, mostly polyhedral with scattered receptor cells. At L3 the geometrically shaped cells have been replaced by elongated cells which represent a transition to the typical vomeronasal receptor epithelium. Only at L5 the medial epithelium for first time shows the typical morphology of a full developed sensory epithelium in a crescent-shaped lumen. At levels L7-L9, the VD has reduced its diameter, its shape is again circular, and three quarters of the epithelial lining of the duct are of neuroreceptor nature, with a clear distinction between neuroreceptor cells, in the basal part and more apical sustentacular cells with dense nuclei. This cellular organization is preserved until the very end of the VD, although there is a remarkable increase in the development of the parenchyma and the size of the VD at L10, which give way suddenly, at L12-13 to the abrupt end of the VD. RE is more uniform. Rostrally squamous, soon is replaced by two layers of rounded or oval cells with scattered superficial cells with translucent nuclei which remain until L4. At L5 still a few polyhedral cells with translucent cytoplasm are seen in its apical zone but the features are mostly typical of a RE. This pattern will be preserved until the caudal extremity of the VD, and even the earliest epithelial invaginations that, like gloved fingers, give origin to the VD are lined exclusively with respiratory epithelium.

The remarkable changes observed along the whole length of the organ in the shape of the lumen have not been previously described in other ruminants, since in the case of the cow, Adams (1986) describes the crescent shape as the predominant one, except for the anterior part of the organ. His description of the rostral part as a cuboidal stratified epithelium of cells with lightly stained cytoplasm coincides what we have observed in the dama gazelle. Likewise, his description as well as that of other authors in cows (Taniguchi & Mikami, 1985; Salazar et al., 2008; Jang et al., 2021) regarding the organization the neuroepithelium in basal, neuroreceptor and sustentacular cells, and the organization of the respiratory epithelium coincides with that observed by us. This is as well comparable to that observed in other Bovinae such as goat (Park et al., 2013), sheep (Kratzing, 1971; Salazar et al., 1998, 2000; Salazar, Lombardero, Alemañ, et al., 2003; Salazar et al., 2007; Ibrahim et al., 2013; Ibrahim, 2018) and water buffalo (Emam et al., 2016), but also with the other ruminants studied, such as giraffe (Kondoh, Nakamura, et al., 2017), reindeer (Bertmar, 1981), elk (Vedin et al., 2010) and Korean roe deer (Park et al., 2014a), which shows that we are dealing with a widely conserved pattern in ruminant evolution.

#### 4.2.2 The vascular pump

The vomeronasal pump is a unique characteristic of the vomeronasal organ found in mammals. This pump is responsible for moving pheromone molecules from the external environment to the sensory cells within the vomeronasal organ, enhancing its chemosensory capabilities. The pumping mechanism is powered by vasomotor movements, which can suck stimulus substances into the vomeronasal organ as well actively expulses the contents of the VD. These mechanisms are activated by fibres running in the nasopalatine nerve which cause constriction of blood vessels within the VNO capsule (Meredith & O’Connell, 1979). After the constriction, the volume of blood in the cavernous tissue is reduced, creating a pressure differential that expands the VNO lumen and draws in fluid from the region around the duct opening (Meredith et al., 1980). Stimuli enter the vomeronasal organ in solution via the mucus stream that flows past the vomeronasal duct from the incisive duct. Effective vomeronasal stimuli should be mucus- soluble chemicals or substances that become mucus-soluble when bound to carrier molecules that may be secreted by vomeronasal glands (Krishna et al., 1995).

The remarkable development of the vascular component of the parenchyma of the dama gazelle is a feature already present in the rostral levels, in which the lateral part of the parenchyma is completely occupied by blood vessels. When moving caudally, the vessels extend towards the dorsal and ventral areas, and finally, in the caudal sections, they occupy the entire parenchyma, as we have observed both in transverse and horizontal sections. This fact is common to ruminant species in which the vascular element has been more studied (Salazar et al., 1998, 2008; Vedin et al., 2010). The presence of a more developed vasculature in the caudal extremity of the VNO is hypothetically necessary for the creation of a powerful negative pressure in this caudal area, because as we have observed in the dama gazelle, a fact not previously described in other species to our knowledge, the vomeronasal neuroepithelium remains nearly until the caudal interdigitations of the vomeronasal duct, which arise directly from the vomeronasal glands in that region. To access the receptors located in this caudal area, a strong negative pressure on the VD at this level is therefore necessary. Access will be determined not only by the action of the pump but also by the physicochemical properties of the molecules. It seems logical to assume that a gradient of dispersion of substances is established along the duct, with only the substances more soluble in the glandular secretion reaching the caudal extremity and the less soluble ones remaining in the anterior part. It seems consistent to hypothesize that the specificity of vomeronasal receptors along the VD is adapted to the differential migration of the mixture of substances along the duct, in the same way as the compounds are eluted in a specific fashion along a chromatography column.

#### 4.2.3 Vomeronasal glands

The vomeronasal epithelia are coated with a mucous fluid produced by glands in the VNO, which is necessary for the neuroepithelium to detect substances (Sankarganesh et al., 2022). As it contains various enzymes that can transform the structure of molecules and alter detection by the receptor neurons the mucus influences olfactory discrimination (Nagashima & Touhara, 2010). Moreover, the mucous fluid contains a number of enzymes with antibacterial and antioxidative properties (Ding & Xie, 2015).

The properties of the mucous fluids differ among the species (Kondoh et al., 2020) and they have been usually characterized employing the Alcian blue (AB) stain that identifies the acidic polysaccharides synthesized and secreted by the glands, and the periodic acid-Schiff (PAS) stain that detects both acidic and neutral polysaccharides. Initial studies on the VNO glands concluded that they secreted only neutral compounds in all species examined (Loo & Kanagasunteram, 1972; Cuschieri & Bannister, 1974; Oikawa et al., 1993, 1994; Salazar et al., 1996). Their results reflected the positivity of the glandular tissue to the PAS marker, while AB produced a negative result. However, in 1997, Salazar and his collaborators showed for the first time AB-positive vomeronasal glands. These findings were found in the VNO of cow and horse, which also showed PAS-positive glands (Salazar, Quinteiro, et al., 1997). Subsequently, several authors obtained the same results by finding both types of VG in all the ruminants studied: cow (Salazar et al., 2008; Jang et al., 2021), sheep (Salazar et al., 2000, 2007), goat (Yang et al., 2021), silka deer (Kondoh et al., 2020), giraffe (Kondoh, Nakamura, et al., 2017), and goitered gazelle (Abood & Hussein, 2018). The degree of development of the glands and the intensity of the response to both stainings, PAS and AB, was similar for all species.

Our observations in the dama gazelle show a clear distinction between the two kinds of secretions. PAS produced a generalized staining of the glandular component from the rostral level L3 to the caudal end of the VNO, whereas Alcian blue only stained the central part of the VNO (level L6) and the staining was restricted to a small group of glands in the dorsal part of the parenchyma. This result confirms that to establish comparison with other group of animals, it is advisable performing studies throughout the entire VNO to avoid making mistakes when extrapolating the information obtained in only a particular area. For instance, this may explain why an initial study in musk shrews did not find AB positive VG (Oikawa et al., 1993); however, more recently AB-positive glands were identified in that species (Kondoh et al., 2020).

#### 4.2.4 Innervation

Although the distribution of the two main components of the innervation of the VNO, the unmyelinated fibers of the vomeronasal nerve and the myelinated fibers of the caudal nasal nerve, have received less attention in studies of the VNO of ruminants, the observations made in the gazelle dama are comparable to those described in other species (Salazar, Quinteiro, et al., 1997; Salazar et al., 2007; Vedin et al., 2010). The serial histological study, however, shows a remarkable feature; already from the most rostral levels (L3) a sensory innervation is present in the medial part of the parenchyma, which means that although the structure of the neuroepithelium at the most rostral levels is still far from the layered structure and the density of receptor cells observed at the central levels, it already possesses a significant sensory innervation.

### 4.3. Immunohistochemical study

Given the remarkable variations observed in the structure of the vomeronasal duct and its epithelial lining, the expression of the markers studied immunohistochemically was not performed only in the central part of the VNO but was extended to the entire length of the VNO.

To characterize the expression of the two major families of vomeronasal receptors, V1R and V2R, respectively (Dulac & Axel, 1995; Ryba & Tirindelli, 1997), specific antibodies against the α subunit of Gi2 and Go proteins are particularly useful. The investigation of G protein expression and distribution in the goat VNS (Takigami, 2000) revealed, for the first time, that the Go pathway had disappeared in some species, a finding that was subsequently confirmed in numerous mammals, including Laurasiatheria and Primates (Suárez et al., 2011). However, there have been observed discrepancies between gene sequencing and immunohistochemical studies, mainly in the Order Carnivora, where Dennis et al. (2003) in the dog and Ortiz-Leal et al. (2020) in the fox observed immunopositive labelling in the vomeronasal neurosensory epithelium using both anti- Gαi2 and anti-Gαo antibodies. This contrasts with the apparent absence of functional V2R genes in the genome of both species (Young & Trask, 2007; Kukekova et al., 2018). However, new sequencing studies that are more focused on olfactory and vomeronasal genes are likely to result in the identification of additional genes that can be added to the repertory of vomeronasal genes identified using a whole-genome approach. In fact, recently Kondoh et al. (2022) detected the presence of functional V2R genes in the genome of cattle, goats, sheep and pigs by genome assembly.

These authors limited their study of Gαi2 and Gαo expression at the AOB, the first integrative center in the brain of the VNS, without performing immunohistochemistry it in the VNO itself. Our study in the dama gazelle may shed some light on this question, as we have observed immunopositivity to both markers Gαi2 and Gαo in the vomeronasal epithelium. Moreover, by performing a serial immunohistochemical study along the entire vomeronasal duct, we have been able to confirm that the expression of both markers is not constant in the SE, since a significant part of it is not immunopositive to either protein. In particular, the large segment extending from L9 to L13, despite having a well characterized neuroepithelium, it lacks immunolabelling for both proteins. This implies that the type of receptor expressed in this caudal third of the VD does not correspond to either V1R or V2R, suggesting that the molecules detected by these receptors do not reach the caudal end of the VD. It follows that only compounds highly soluble in the vomeronasal mucus and probably of low molecular weight can be detected in the caudal VD and probably for another kind of receptors yet not described. The absence of vomeronasal receptor expression in such a large stretch of the VNO could explain why the previous studies of G-protein expression in the ruminant VNO have failed to found

Gαo immunopositivity, as it has been the case in goat (Takigami, 2000), sheep (Salazar et al., 2007), and silka deer (Matsubara et al., 2019). The choice of a single level, usually the central one, for these studies might have conditioned the results obtained.

In the discussion of Gαo/V2R expression, it is worth noting that Kondoh et al. (2022) in their study of the AOB of the cow did not find immunopositivity to Gαo, nor was it found in the AOB of the dog (Salazar et al., 2013) or fox (Ortiz-Leal, Torres, Villamayor, et al., 2022), but this does not exclude the expression of V2R in the VNO, as it is plausible that information from these receptors may project to other areas of the olfactory bulb, not necessarily the AOB. In fact, recently, it has been found in the olfactory bulb of the fox, that the expression of Gαo linked to the vomeronasal nerves projects not to the AOB, but the transition zone located between the main and the accessory olfactory bulb, area known as the olfactory limbus (Ortiz-Leal et al., 2023).

The neuronal maturity of the VNO neuroepithelium in the dama gazelle was analysed with anti-OMP, widely used as a molecular marker for olfactory neurons in both the olfactory mucosa and the VNO in different species (Smith, Dennis, et al., 2011; Sasuga et al., 2013; Rodewald et al., 2016). The functional properties of OMP are still a matter of debate. It has been shown to play a role in the maturation of olfactory neurons and for the level of selectivity in stimulus-response (Buiakova et al., 1996).

In the dama gazelle anti-OMP produced immunopositive labelling in the vomeronasal sensory epithelium from the rostral levels, but it did not stain the caudal levels L10-L13 as it also happened with the G-proteins. Anti-OMP stained cell bodies more intensely than dendrites in OMP-positive receptor cells. This feature was also found in sheep (Ibrahim, 2018), cow (Jang et al., 2021), and roe deer (Shin et al., 2017), however, goat showed the reverse pattern (Park et al., 2013; Yang et al., 2021). Anti-OMP immunolabelling in the RE shows variations between the different ruminants. While the ewe shows no positive OMP marking in its RE (Salazar et al., 2007), the roe deer has ciliated and basal cells OMP positive (Shin et al., 2017). In contrast, the dama gazelle shows an intermediate pattern, with diffuse labelling and only a few ciliated OMP-positive cells in its RE. Likewise, both the cow and Korean black goat show this intermediate pattern (Park et al., 2013; Yang et al., 2021).

To gain a better understanding of the neurochemical properties of the vomeronasal neuroepithelium, we made use of an additional range of neuronal markers. Protein gene product 9.5 (PGP 9.5) is a neuronal biomarker expressed in sensory paraneurons that plays a vital role in ubiquitin regulation. It has been found expressed in olfactory receptor cells, taste bud cells and vestibular hair cells (Iwanaga et al., 1992; Johnson et al., 1994). The results obtained by us in gazelle dama are consistent with those observed in sheep (Ibrahim, 2018), cows (Jang et al., 2021), Asian roe deer (Park et al., 2014a) and Korean black goats (Park et al., 2013). PGP 9.5-positive cells show strong immunoreactivity in their soma, dendrites, and their axons which extend through the lamina. Additionally, some PGP 9.5-positive ciliated cells are observed in the RE, as is the case in all ruminates mentioned except sheep.

According to the findings of our investigation conducted along the VNO rostrocaudal axis, PGP 9.5 expression can be found in the caudal sensory epithelium zone (L10-L13), even though neither G proteins nor OMP are expressed there. On one hand, this provides evidence that the histologically reported neuroepithelial nature of this long stretch of medial epithelium is correct. Furthermore, it suggests that the sort of receptors that are expressed are of a different kind than those belonging to the V1R and V2R families.

Neuron-specific enolase (NE), a brain-specific isozyme of the glycolytic enzyme enolase, is characterized by its consistent occurrence in the cytoplasm of mature neurons (Iwanaga et al., 1989), including sensory neurons. To our knowledge there are no immunohistochemical studies performed with this marker in ruminants. Our observations in the dama gazelle not only confirm the immunolabelling obtained in a wide variety of mammalian orders, including rodents and even humans. (Takami et al., 1993; Wang et al., 1994), but also extend the immunopositivity to the entire sensory epithelium, confirming the results obtained with PGP 9.5 regarding the neurosensorial nature of the caudal third of the VD.

Calcium binding proteins are specific markers of vomeronasal receptor cells, in which they play a role in the modulation of calcium signalling and therefore in the transmission of sensory information. Although their presence has been identified in the VNO of a wide range of mammalian groups including Rodentia (Kishimoto et al., 1993; Jia & Halpern, 2003; Torres et al., 2020), Marsupialia (Jia & Halpern, 2004; Torres et al., 2022), Canidae (Ortiz-Leal et al., 2020), Soricidae (Malz et al., 2000) and Hominidae (Johnson et al., 1994), there is however no information available on this group of proteins in the VNO of ruminants. By studying four of them - calbindin, calretinin, parvoalbumin and secretagogin - we have been able to fill an important gap in the study of the neurochemistry of the vomeronasal organ in such a large group of mammals. In the specific case of secretagogin, this is the first description of the expression of this marker, recently discovered (Wagner et al., 2000) in the mammalian VNO. Of the four markers used, calbidin and calretinin have given the best results in characterizing the structural elements that form the neuroreceptor epithelium of the VNO, since the expression of secretagogin and parvoalbumin was common to both sensory and respiratory epithelia. As it happens in the dama gazelle, anti-parvalbumin stained in the mouse the VNO moderately and less intensely than CB or CR (Kishimoto et al., 1993).

Both antibodies against CB and CR stained a subpopulation of vomeronasal receptor neurons in the dama gazelle VD. Somata of the immunoreactive neurons were located in the basal and middle region of the sensory epithelium of the VNO, which contrasts with the apical position of CB somata in the rat VNO (Jia & Halpern, 2003). Thick apical dendrites, piriform somata, and axons were clearly immunostained with both markers in the dama gazelle. In addition, CB-and CR- immunoreactive axons could be found in the nerve bundles in the lamina propria and in the vomeronasal nerves. This immunolabeling pattern first started to become visible in the SE at rostral levels L3-L4 in a gradual way, and then generalized to the entire SE from level L5 to L9. As in the case of G-proteins and OMP expression, the expression of calcium-binding proteins did not occur in the caudal zone of the sensory epithelium.

The immunohistochemical study was completed with a specific characterization of the epithelial tissue using the anti-cytokeratin 8 antibody, a type II intermediate filament protein (Franke et al., 1981). In the dama gazelle anti-CYK8 identified neuroreceptor cells and sustentacular cells across the sensory and respiratory epithelia in a broad manner. Only in the anterior levels of the VD, anti-CYK8 did not produced an homogeneous labelling, unstaining the apical polygonal cells, whose functional significance remains unknown as we could not find any reference to this cell type in the VNO literature. Most of the investigations of cytokeratins in the VNO have been done using cytokeratin cocktails. Dennis et al. (2003) found in dogs that the immunolabelling was restricted to the basal cells, whereas Poran (1998) found in the opossum a more diffuse pattern, similar to the one we observed in the dama gazelle, albeit with greater intensity in the RE than in the SE.

### 4.4 Lectin histochemical study

Lectins are a type of agglutinin which is capable of binding to certain glycoconjugates that are found on the surface of cells (Brooks, 2023). Since the luminal surface of the vomeronasal duct is covered by a layer of glycocalyx, which is a layer of carbohydrates and proteins, the utilization of these molecules is particularly helpful in the investigation of the vomeronasal system (Plendl & Sinowatz, 1998). Moreover, lectins are used to identify and mark the sensory cells, the respiratory epithelium and the glandular component of the VNO (Halpern et al., 1998). If we limit our focus to the nervous tissue, it is remarkable how certain lectins are unique to the vomeronasal pathway. These lectins recognize the overall neuroreceptor cell, not only at the level of its dendrites and somas, but also of its axons, all the way up to its terminal projections in the accessory olfactory bulb (Salazar et al., 1994; Kondoh, Kamikawa, et al., 2017; Kondoh et al., 2018; Chun et al., 2023). However, any study of the vomeronasal system based on the use of lectins must take into account that the labelling patterns are highly specific, with notable differences even between species as phylogenetically close as the rat (Salazar & Sánchez Quinteiro, 1998) and the mouse (Salazar et al., 2001), and extrapolations should be avoided.

In our investigation on the VNO of the dama gazelle, we have employed five lectins to reveal the glycoconjugates binding patterns in both the vomeronasal sensory and respiratory epithelia, the vomeronasal nerves and glands. All the lectins employed stained both the vomeronasal glands (VG) and the muco-microvillar complex, which covers the luminal surface of both vomeronasal epithelia. In addition, each lectin specifically stained different structures of the VNO and exhibited its own localisation patterns and labelling intensity. The lectin SBA (*Glycine max*) produced intense labelling in the glandular tissue all through the VNO, from its rostral extremity. Similarly, VVA (*Vicia villosa*) stained the VG in the central part of the VNO, but much less intensely than SBA. Neither SBA nor VVA stained the VD epithelia nor the vomeronasal nerve axons. LEA (*Lycopersicon esculentum*) and STA (*Solanum tuberosum*) lectins strongly labelled the vomeronasal epithelium, vomeronasal glands (VG) and vomeronasal nerves, while the caudal nasal nerve remained unstained. As for the lectin UEA (*Ulex europaeus*), its labelling is concentrated in the vomeronasal epithelium and VG, however, it did not stain the vomeronasal nerves in the parenchyma of the VNO. In contrast to the proteins detected in the immunohistochemical study, there were no noticeable differences in the lectin-histochemical labelling pattern along the rostro-caudal axis of the VNO.

These results show differences and similarities with the labelling obtained by these lectins in the VNO of other ruminant mammalian species. For this reason, our findings are related below to existing information on sheep (*Ovis orientalis aries*) (Salazar et al., 2000, 2007; Ibrahim et al., 2013), cow (*Bos taurus coreanae*) (Jang et al., 2021), goat (*Capra aegagrus hircus*) (Park et al., 2013; Yang et al., 2021) and Asian roe deer (*Capreolus pygargus*) (Park et al., 2014b; Shin et al., 2017).

SBA produced analogous labelling in dama gazelle and sheep (Ibrahim, 2018) and, as in these species only the VG and the luminal rim of the vomeronasal epithelium were stained. Additionally, the same marker moderately stained the vomeronasal SE in ruminant species such as cow (Jang et al., 2021), goat (Park et al., 2013; Yang et al., 2021) and roe deer (Park et al., 2014b). In cows it stains as well moderately the vomeronasal nerves (Jang et al., 2021) VVA lectin only marked the superficial border of the epithelium and the VG in the dama gazelle, although less intensely than SBA. In contrast, the labelling of this lectin extends moderately to the vomeronasal epithelium and NVN in the cow (Jang et al., 2021), and to a lesser extent, in the sheep (Ibrahim et al., 2013).

LEA stained numerous vomeronasal structures in the dama gazelle: the vomeronasal epithelial rim, the vomeronasal epithelium, the VGs and the NVNs. In the cow, sheep, and roedeer, LEA stains the same structures in a similar fashion (Salazar et al., 2000; Ibrahim et al., 2013; Shin et al., 2017; Jang et al., 2021). Interestingly, STA follows exactly the same pattern as LEA. It marks the luminal rim, vomeronasal epithelium, VGs and NVNs in dama gazelle, roe deer and cow (Shin et al., 2017; Jang et al., 2021). Finally, the UEA lectin marked three vomeronasal structures in the VNO of the dama gazelle: the luminal rim, the vomeronasal epithelium and the VGs. This pattern is repeated in sheep, goat, roe deer and cow although faintly (Salazar et al., 2000; Ibrahim et al., 2013; Park et al., 2013; Shin et al., 2017; Jang et al., 2021). In cow, UEA also stains NVNs (Jang et al., 2021).

In summary, our study has contributed to fill the existing gap in the knowledge of the vomeronasal organ in wild ruminant species, which are a group of animals for which little information is available on the histological and neurochemical organization of their vomeronasal system. By investigating the dama gazelle vomeronasal organ, we have gained valuable insights into the structure and function of this critical sensory structure in this endangered species. Moreover, as an threatened species, the information obtained from our study can be used to design and implement programs that utilize pheromones to enhance breeding success and increase genetic diversity in captive populations. Such programs may be critical for the survival of the species, and our work has provided a solid morphological foundation for future research on this topic.

Finally, our study has also made a significant contribution to the general study of the vomeronasal organ. By conducting a comprehensive histological and immunohistochemical study of the organ, we have identified notable differences in the organization of the vomeronasal duct and the expression of neuromarkers along the rostrocaudal axis of the organ. These findings highlight the importance of considering such differences when conducting future studies of the vomeronasal organ in other species. Overall, our study underscores the critical role played by the vomeronasal organ in mediating social and reproductive behaviors in the dama gazelle and highlights the need for further research to better understand this unique sensory structure in other wild ruminant species.

## AUTHORS CONTRIBUTION

M.V.T., I.O.L., P.S.Q. designed the research., M.V.T., I.O.L., P.S.Q., A.F., J.L.R. performed the work, M.V.T., I.O.L., P.S.Q. analysed and discussed the results and wrote the paper.

## COMPLIANCE OF ETHICAL STANDARDS

### Ethical approval

All the animals employed in this study dead by natural causes.

### Informed consent

No human subject was used in this study.

## ACKNOWLEDGEMENTS

The authors thank MARCELLE NATUREZA park (Outeiro de Rei, Spain) for providing the animal employed in this study. The authors would like to express their gratitude to Dr. Ludwig Wagner (University of Vienna) for kindly donating the antibody against secretagogin.

